# AMPK activation by metformin promotes survival of dormant ER+ breast cancer cells

**DOI:** 10.1101/2020.01.21.914382

**Authors:** Riley A. Hampsch, Jason D. Wells, Nicole A. Traphagen, Charlotte F. McCleery, Jennifer L. Fields, Kevin Shee, Lloye M. Dillon, Darcy B. Pooler, Lionel D. Lewis, Eugene Demidenko, Yina H. Huang, Jonathan D. Marotti, William B. Kinlaw, Todd W. Miller

**Affiliations:** Department of Molecular & Systems Biology, Norris Cotton Cancer Center, Geisel School of Medicine at Dartmouth, Lebanon, NH; Department of Microbiology & Immunology, Norris Cotton Cancer Center, Geisel School of Medicine at Dartmouth, Lebanon, NH; Department of Medicine, Norris Cotton Cancer Center, Geisel School of Medicine at Dartmouth, Lebanon, NH; Department of Community & Family Medicine, Norris Cotton Cancer Center, Geisel School of Medicine at Dartmouth, Lebanon, NH; Department of Pathology, Norris Cotton Cancer Center, Geisel School of Medicine at Dartmouth, Lebanon, NH; Department of Comprehensive Breast Program, Norris Cotton Cancer Center, Geisel School of Medicine at Dartmouth, Lebanon, NH

**Author notes:** To whom correspondence should be addressed. **Corresponding author:** Todd W. Miller, Dartmouth-Hitchcock Medical Center, One Medical Center Dr., HB-7936, Lebanon, NH 03756.

**Keywords:** breast cancer, drug resistance, anti-estrogen, dormancy

## Abstract

**Purpose:** Despite adjuvant anti-estrogen therapy for patients with estrogen receptor alpha (ER)-positive breast cancer, dormant residual disease can persist for years and eventually cause tumor recurrence. We sought to deduce mechanisms underlying the persistence of dormant cancer cells to identify therapeutic strategies.

**Experimental Design:** Mimicking the aromatase inhibitor-induced depletion of estrogen levels used to treat patients, we developed preclinical models of dormancy in ER+ breast cancer induced by estrogen withdrawal in mice. We analyzed tumor xenografts and cultured cancer cells for molecular and cellular responses to estrogen withdrawal and drug treatments. Publicly available clinical breast tumor gene expression datasets were analyzed for responses to neoadjuvant anti-estrogen therapy.

**Results:** Dormant breast cancer cells exhibited upregulated 5’ adenosine monophosphate-activated protein kinase (AMPK) levels and activity, and upregulated fatty acid oxidation. While the anti-diabetes AMPK-activating drug metformin slowed the estrogen-driven growth of cells and tumors, metformin promoted the persistence of estrogen-deprived cells and tumors through increased mitochondrial respiration driven by fatty acid oxidation. Pharmacologic or genetic inhibition of AMPK or fatty acid oxidation promoted clearance of dormant residual disease, while dietary fat increased tumor cell survival.

**Conclusions:** AMPK has context-dependent effects in cancer, cautioning against the widespread use of an AMPK activator across disease settings. The development of therapeutics targeting fat metabolism is warranted in ER+ breast cancer.

**Statement of Translational Relevance:** Dormant cancer cells that survive adjuvant therapy can ultimately give rise to recurrent/advanced tumors that frequently develop resistance to all approved therapies. Patients with early-stage estrogen receptor alpha (ER)-positive breast cancer are typically treated with surgical resection followed by ≥5 years of adjuvant anti-estrogen therapy that neutralizes ER and suppresses, but often does not eliminate, tumor-initiating cells. Estrogen withdrawal, which mimics aromatase inhibitor therapy, induced activation of the metabolic sensor 5’ adenosine monophosphate-activated protein kinase (AMPK) and upregulated fatty acid oxidation (FAO) in preclinical models. Treatment with the anti-diabetes AMPK-activating drug metformin or high dietary fat intake promoted survival of dormant ER+ breast cancer cells, while anti-anginal drugs that inhibit FAO induced clearance of dormant tumor cells. These findings caution against using AMPK modulators with anti-estrogens in patients with ER+ breast cancer, and warrant testing of FAO inhibitors as anti-cancer agents in combination with anti-estrogens.

## Introduction

Estrogen receptor α-positive (ER+) breast cancer (BC) is commonly treated with anti-estrogens that target tumor-driving ER activity by either inhibiting ER directly (*e.g.*, tamoxifen), or reducing estrogen levels through suppression of biosynthesis [*e.g*., aromatase inhibitors (AIs), such as letrozole]. While adjuvant anti-estrogen therapies have shown clinical benefit for the treatment of ER+ BC, ∼30% of patients eventually experience disease recurrence. Recurrent disease is ultimately fatal in a majority of cases, as cancers can rapidly become refractory to standard therapies. The bulk of these recurrence events occur after the standard five-year adjuvant anti-estrogen treatment regimen (termed “late recurrence”) (1–3). This lengthy time to recurrence suggests that ER+ BC cells undergo extensive periods of dormancy, defined as disease that is undetectable by conventional clinical follow-up techniques (*e.g.*, PET/CT). Indeed, circulating tumor cells (CTCs) in blood and disseminated tumor cells (DTCs) as well as micrometastases in bone marrow are detectable in “disease-free” BC patients after years of adjuvant anti-estrogen therapy (4, 5). Thus, there is a need for development of more effective adjuvant therapeutics to either maintain this residual disease in a dormant asymptomatic state, or to eradicate it. Such therapeutic development has been hampered by a limited understanding of the biology underlying dormancy.

Several groups have developed preclinical models of cancer cell dormancy and identified mechanisms driving emergence from dormancy (6, 7), with the intent to develop therapeutics to suppress tumor outgrowth. We postulated that understanding mechanisms that allow cancers cells to survive/persist in a dormant state would lead to curative therapies (*i.e.*, cancer elimination) rather than suppressive therapies (*i.e.*, maintenance of dormancy). Moreover, reported dormancy models often utilize therapy-naïve contexts, while more clinically relevant models would include (neo)adjuvant therapies. Additionally, there is a dearth of *in vivo* models of dormancy in ER+ BC, a cancer in which dormancy is exceedingly clinically relevant due to the long latency of recurrences and the steady lifetime risk of recurrence (8). Herein, we developed *in vivo* models of estrogen withdrawal (EW)-induced dormant ER+ BC to identify mechanisms of BC cell survival/persistence, reflecting the clinical scenario of patients on (neo)adjuvant AI therapy.

## Materials and Methods

### Cell culture and RNA interference

Parental cell lines (ATCC) were cultured in DMEM with 10% FBS. For hormone deprivation (HD) experiments, cells were cultured in phenol-red free DMEM containing 10% dextran/charcoal-stripped FBS ± 1 nM E2. Cells were stably transfected with lentiviral vectors encoding luciferase, GFP, shRNA targeting AMPKα1/PRKAA1 or non-targeting control. Cells were transiently transfected with siRNA targeting AMPKα1/*PRKAA1*, AMPKα2/*PRKAA2*, and/or non-silencing control.

### Tumor growth studies

Animal studies were approved by the Dartmouth College IACUC. Female NOD-scid/IL2Rγ^−/−^ (NSG) mice or *Foxn1^nu^*/*Foxn1^nu^* (nude; JNu) mice (3-4 wk) were ovariectomized (ovx) and orthotopically injected bilaterally with MCF-7, HCC-1428, HCC-1500, or MDA-MB-415 luciferase/GFP-expressing cells, or implanted with fragments of HCI-017 patient-derived xenograft (PDX; a gift from Alana Welm, Univ. of Utah). Mice were simultaneously implanted s.c. with a 17β-estradiol (E2) pellet as indicated. Tumor dimensions were measured twice weekly using calipers (volume = [length^2^ × width]/2). Once tumors reached ∼400 mm^3^, E2 pellets were removed [*i.e.*, estrogen withdrawal (EW)] and mice were randomized to treatments. Vehicle or metformin (1 mg/mL) were administered via drinking water containing 4% sucrose. Once tumors completely regressed (*i.e*., not palpable) following EW, residual tumor burden was monitored by bioluminescence imaging. For drugs tested against dormant residual disease: tumor-bearing mice were treated with EW for 60-90 d to induce dormancy; baseline bioluminescence was measured on two consecutive days and averaged; animals were treated as indicated and bioluminescence was serially measured. For molecular analysis, tumors were harvested at the indicated time points, fixed in formalin, and paraffin-embedded (FFPE).

### Statistical analyses

Cell growth data and immunohistochemical (IHC) scores were analyzed by *t*-test (for two-group experiments), or ANOVA (for experiments with more than two groups) followed by Bonferroni multiple comparison-adjusted posthoc testing between groups. Mathematical models used to analyze tumor volumes and bioluminescence are described in Supplemental Methods.

**Additional details are provided in Supplemental Methods**

## Results

### A subpopulation of ER+ BC cells survives extended periods of estrogen withdrawal in a dormant state

Laboratory mice have low circulating levels of estrogens, and ovx further reduces systemic estrogen levels (9). Human ER+/HER2− BC cells commonly require estrogen supplementation to form tumors in immunodeficient mice. Ovx NSG mice injected subcutaneously (s.c.) with MCF-7 cells and supplemented with E2 formed visible tumors within 3 wk. Mice injected with MCF-7 cells without E2 supplementation did not form palpable tumors within 10 wk; however, E2 supplementation at the 10-wk time point induced tumor formation (Fig. S1). This indicates that a proportion of MCF-7 cells survive extended estrogen withdrawal (EW) in a non-palpable, dormant state while retaining tumorigenicity.

We then developed a platform for the preclinical study of dormancy in ER+ BC (Fig. 1A). ER+/HER2− BC cell lines stably expressing luciferase (MCF-7, HCC-1428, HCC-1500, or MDA-MB-415) or HCI-017 PDX tumor fragments were orthotopically implanted in the mammary fat pads of ovx immunodeficient (NSG or nude) mice, and mice were supplemented with E2. E2-driven tumors were grown to ∼400 mm^3^ to allow for establishment of tumor microenvironment and vasculature. E2 was then withdrawn to mimic the EW induced by AI therapy in patients. EW caused rapid tumor regression to a non-palpable state in all (MCF-7, HCC-1500, and HCI-017) or most (HCC-1428 and MDA-MB-415) cases (Figs. 1B and S2A-E). Serial non-invasive bioluminescence imaging indicated that residual tumor cell burden stabilized after ∼60-90 d of EW and persisted for >1 yr (Fig. 1C/D). For subsequent analyses, we chose 60-90 d of EW as a window to evaluate dormant tumor cells. These models reflect the clinical scenario of minimal residual disease that can give rise to recurrent tumors.

**Fig. 1.**
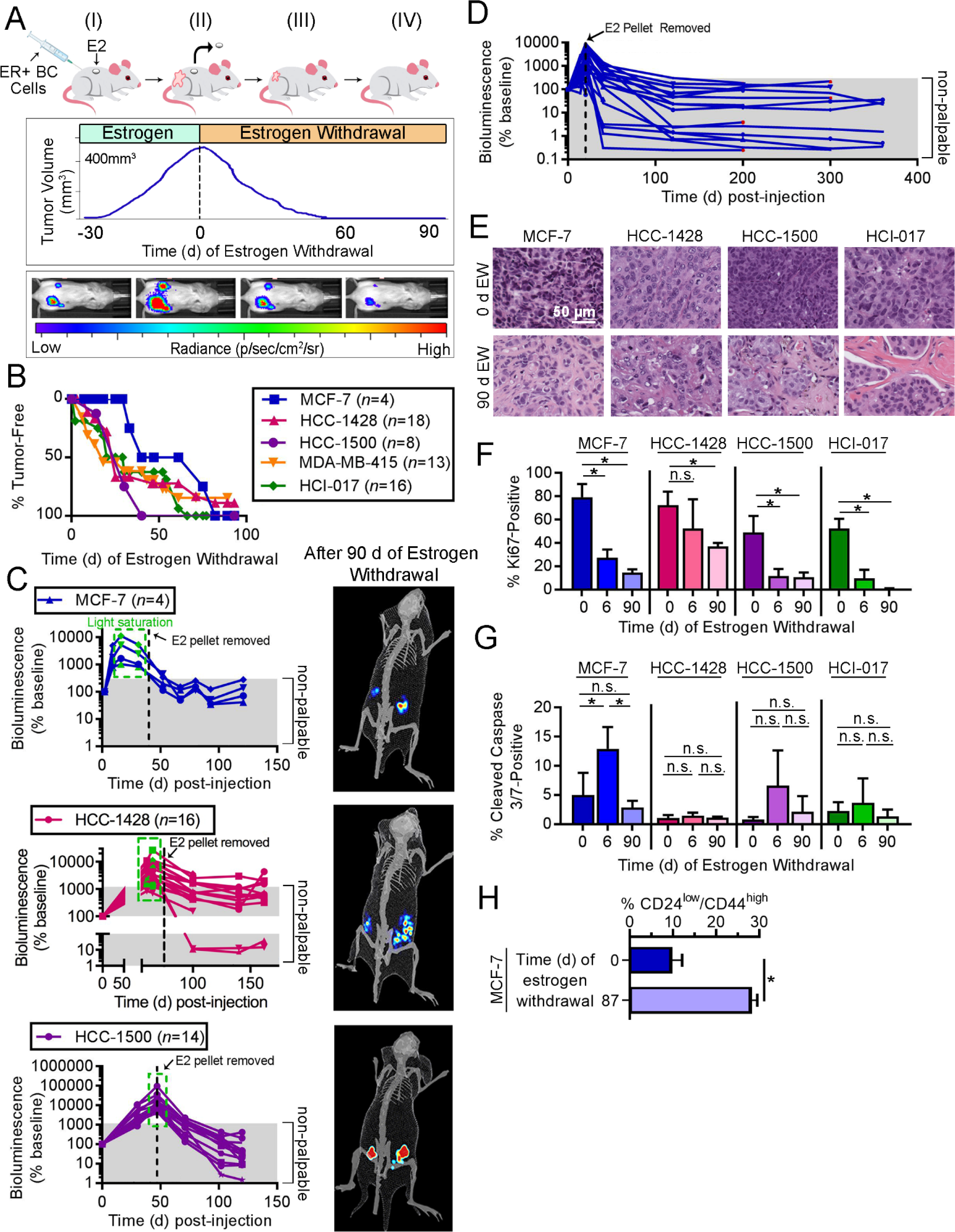
Modeling dormancy in ER+ BC. **(A)** Outline of modeling. (I) Ovx mice are injected orthotopically with luciferase/GFP-labeled ER+ BC cells or a PDX tumor fragment, and supplemented with E2. (II) After 30-60 d, tumors grow to ∼400 mm^3^, and E2 pellets is withdrawn (EW). (III) Tumor regression and residual cell burden are measured using calipers and bioluminescence imaging. (IV) After 60-90 d of EW, bioluminescence signal (reflecting surviving BC cells) is stable. **(B-D)** Ovx mice injected orthotopically with ER+ BC cells or HCI-017 tumor fragments were supplemented with E2, followed by EW and serial tumor volume measurement (B) and/or bioluminescence imaging (C/D). In E2-supplemented mice with visible tumors, bioluminescence signal was sometimes saturating (green boxes). Following EW (dotted black lines), most tumors regressed beyond palpability. Representative exposure-matched bioluminescence images from mice after 90 d of EW are shown. In (D), MCF-7 tumor regrowth occurred in 4/20 cases after ∼200 d of EW (red points); these tumors modeled resistance to EW, and were not analyzed herein. **(E)** Representative H&E-stained images of xenografts after 0 or 90 d of EW. **(F/G)** IHC for Ki67 (F) or cleaved caspase 3/7 (G) was performed on xenografts after 0, 6, or 90 d of EW. **(H)** GFP+ MCF-7 tumors were harvested from ovx mice after 0 or 87 d of EW. Tumors were enzymatically digested, and single-cell suspensions were analyzed by flow cytometry for CD24 and CD44. In (F-H), data are shown as mean of triplicates +SD. *p≤0.05 by Bonferroni multiple comparison-adjusted posthoc test (F/G) or *t*-test (H). n.s.-not significant.

Histological analysis of dormant tumor beds after 90 d of EW revealed involutional-type changes, decreased tumor cellularity, increased stromal fibrosis, and extracellular deposition of basement membrane when compared to E2-driven tumors at baseline (Fig. 1E). Previous investigations of cancer dormancy have led to the characterization of two non-mutually exclusive states: 1) population-level dormancy, where insufficient resources lead to a balance between tumor cell proliferation and death that prevents tumor outgrowth; 2) cellular dormancy, where cells are arrested in G_0_ (10). Proportions of Ki67-positive tumors cells were significantly decreased after 6 d (short-term) and 90 d (long-term) of EW compared to baseline (E2-driven), indicating reduced proliferative rates typically associated with cellular dormancy (Figs. 1F and S3A); these data are consistent with the reduced rates of Ki67-positivity observed in residual human tumors following 3 months of neoadjuvant AI therapy (11). Low proportions of apoptotic cells were detected in tumors acquired at baseline and after 90 d of EW, while significantly increased apoptosis after short-term EW was observed only in MCF-7 tumors (Figs. 1G and S3B). As these dormant tumor cells can persist for >1 yr and only infrequently regrow palpable estrogen-independent tumors (Fig. 1D), population-level dormancy may also be occurring. Dormant tumor cells maintained variable degrees of ER positivity and frequent tumorigenic response to estrogen re-challenge (Figs. S2F-H and S3C-D). Consistent with the variability in ER levels, we found that dormant MCF-7 tumor cells were enriched for the CD24^low^/CD44^high^ cancer stem cell phenotype (Fig. 1H); this reflects the ER-low phenotype of stem cells derived from human ER+ tumors (12). Although DTCs and CTCs are frequently ER- in patients with primary ER+ breast tumors, recurrent tumors and metastases typically have ER concordance with primary tumors (13–15), implying that ER-low cancer cells can give rise to ER+ tumors.

### Dormant ER+ BC cells exhibit upregulated AMPK levels and consequent metabolic activities

RNA isolated from formalin-fixed, paraffin embedded (FFPE) specimens of E2-driven (baseline), acutely EW (6 d following EW), or dormant (90 d following EW) MCF-7 and HCC-1428 xenografts from nude mice was used for transcriptome-wide sequencing of RNA (RNA-seq). Gene set variation analysis (GSVA) of transcriptome expression profiles of dormant tumors compared to baseline revealed enrichment for activation of AMPK, FAO, and lipid metabolism pathways (Figs. 2A and S4). As expected, pathways such as ‘Cell Division’ and ‘E2F Targets’ were also significantly altered in dormant tumors. In a separate experiment with MCF-7 tumors harvested at similar time points from NSG mice, we digitally quantified transcripts from 770 genes using the NanoString PanCancer Pathways platform (Table S1). *PRKAA2* mRNA, which encodes the α2 catalytic subunit of AMPK, was upregulated in dormant tumors, but not acutely EW tumors, compared to baseline in all three experiments (Figs. 2B and S5).

AMPK is a cellular energy sensor that promotes catabolism in response to high AMP:ATP ratios or other stress signals. AMPK has a diverse range of cellular activities including, but not limited to, direct and indirect inhibition of mechanistic target of rapamycin complex 1 (mTORC1), induction of autophagy, induction of mitochondrial biogenesis and function through activation of peroxisome proliferator-activated receptor-γ coactivator-1α (PGC-1α) and subsequent transcriptional signaling, and regulation of fatty acid metabolism through inhibitory phosphorylation of acetyl-CoA carboxylase (ACC) (16). IHC analysis showed increased AMPKα2 levels in dormant MCF-7, HCC-1428, and HCC-1500 tumor cells compared to E2-driven tumors (Figs. 2C and S6). P-ACC_Ser79_, a marker of AMPK stimulation of fatty acid β-oxidation (FAO), was increased in dormant tumor cells compared to E2-driven or acutely EW tumors in 3/4 xenograft models, demonstrating increased AMPK activity in dormant tumor cells (Figs. 2D and S6). In contrast, mTORC1 activity was not suppressed: dormant tumor cells exhibited elevated levels of P-S6, a downstream marker of mTORC1 activity, compared to E2-driven tumors (Fig. S7).

Neoadjuvant treatment with anti-estrogens frequently induces tumor regression but not pathologic complete response in post-menopausal women with early-stage ER+ breast cancer (17). To determine whether residual tumor cells following anti-estrogen therapy exhibit alterations in markers of FAO, we evaluated transcriptional profiles of tumor specimens acquired at baseline and after neoadjuvant treatment from three patient cohorts. We derived a 992-gene expression signature of FAO inhibition using transcriptional profiles from MCF-7 cells treated +/− the CPT1 inhibitor etomoxir, which blocks FAO; this signature was used to generate an ‘etomoxir *t*-statistic’ for each tumor specimen, where the additive inverse is inferred to reflect FAO activation. In the GSE20181 cohort, patients were treated with neoadjuvant AI therapy with letrozole, and tumor specimens were serially obtained at baseline, Day 10-14, and Day 90 (18). Tumors showed transcriptional profiles reflecting increased FAO on Day 90, but not Day 10-14, compared to baseline (Fig. 2E). Similar results were observed in the GSE71791 cohort of patients treated with the anti-estrogen fulvestrant for 28 d (19). The GSE111563 cohort contained patients treated with neoadjuvant letrozole for 4-45 months who experienced >40% decrease in tumor size within 4 mo; tumors were biopsied after 13-120 d of letrozole, and those with tumor progression or an increase in mRNA markers of cell proliferation (in the 13-to-120-day biopsy) were classified as “acquired resistant,” while the remainder were classified as “dormant” (17). “Dormant” tumors showed transcriptional profiles of FAO activation, while “acquired resistant” tumors did not (Fig. 2E). Gene Ontology annotations for 77 of 992 genes in the etomoxir signature included “Cell Proliferation” or “Cell Cycle,” so *t*-statistics were also generated for each tumor using the remaining 915 genes to minimize the contribution of proliferated-related transcripts; similar results were observed (Fig. S8), indicating that tumor cells that persist during anti-estrogen treatment exhibit transcriptional profiles reflecting increased FAO.

**Fig. 2.**
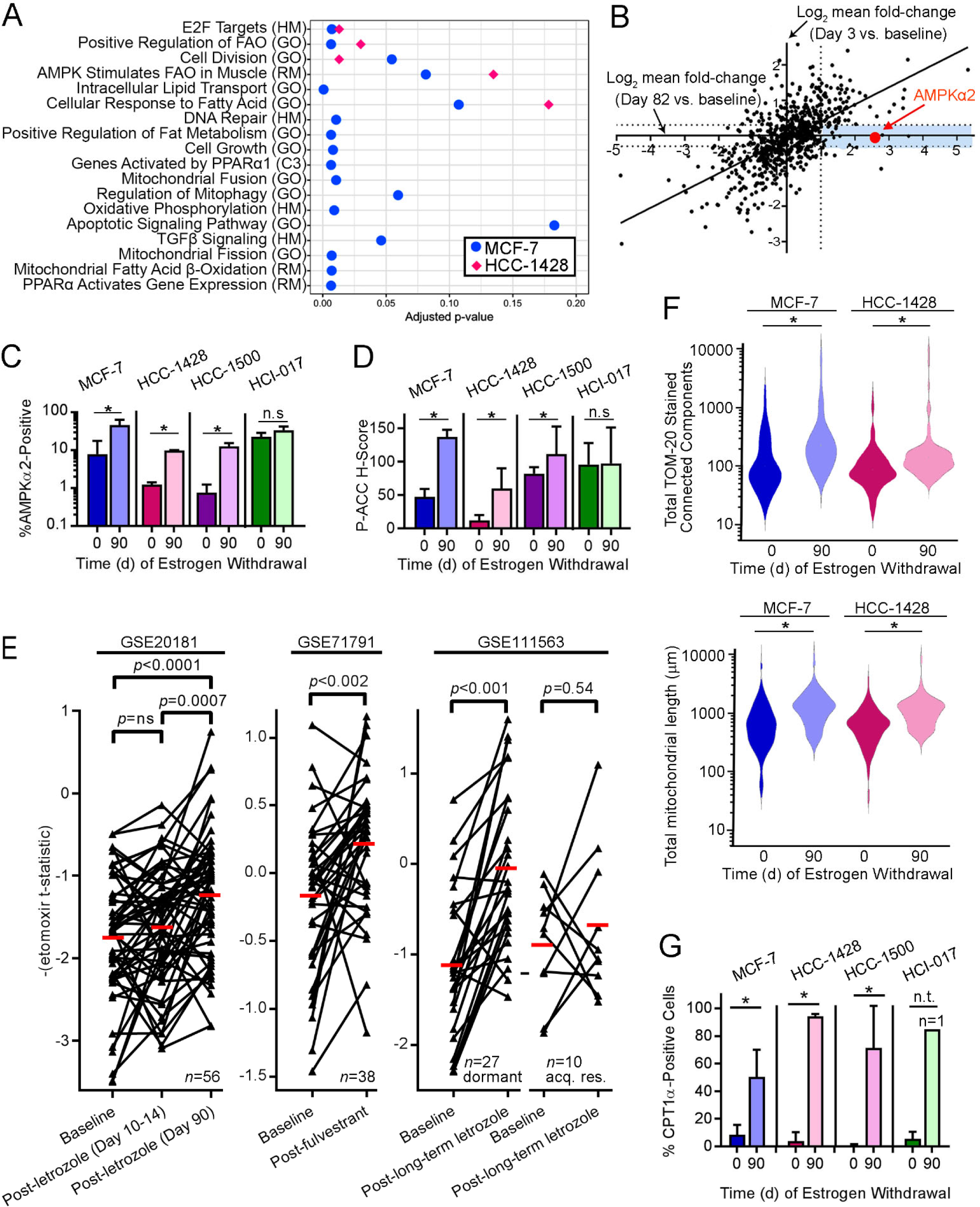
Dormant ER+ breast tumor cells exhibit increased AMPK levels and activity. **(A)** RNA from MCF-7 and HCC-1428 xenografts after 0 or 90 d of EW (*n*=3-4 per time point) was analyzed by RNA-seq. Gene expression changes detected in 90-day tumors vs. baseline were analyzed by unsupervised sample-wise enrichment analysis of metabolic and signaling gene sets using GSVA. **(B)** RNA from MCF-7 xenografts after 0 d (*n*=2), 3 d (*n*=3), or 82 d (*n*=4) of EW was analyzed using the NanoString PanCancer Pathways Panel. Mean fold-change for each gene at 3 d or 82 d relative to baseline was calculated. Blue shading highlights genes with expression increased ≥2-fold in dormant tumor cells (82 d) but not in acutely EW tumors (3 d). **(C/D)** IHC for AMPKα2 and P-ACC_Ser79_. Data are shown as mean of triplicates +SD. \**p*≤0.05 by *t*-test compared to control. n.s.-not significant. **(E)** Human ER+ breast tumor etomoxir *t*-statistics from tumors sampled before and after neoadjuvant anti-estrogen therapy were compared by paired *t*-test. acq. res.-acquired resistant. Red bars indicate mean values. **(F)** Mitochondria were stained with TOM20-AF594 using triplicate tumors. Mitochondrial count (*left*) and length (*right*) per cell (from ≥30 cells/tumor) are shown as violin plot. \**p*≤0.05 by *t*-test. **(G)** IHC of CPT1α. * *t*-test *p*≤0.05. n.t.-not tested (lack of replicates).

If AMPK-directed energy metabolism is increased in dormant tumor cells, then mitochondrial content, cellular respiration, and FAO enzymes may be increased in parallel. Indeed, mitochondria-encoded mRNA levels were increased in dormant MCF-7 tumors compared to E2-driven and acutely EW tumors (Fig. S9A). Morphological assessment revealed increased numbers of mitochondria and mitochondrial elongation in dormant tumors (Figs. 2F and S9B/C), indicative of increased levels and activity of mitochondria (20). Dormant tumors showed increased levels of CTP1α, which is the rate-limiting enzyme in mitochondrial FAO, compared to E2-driven tumors (Figs. 2G and S9D). These data collectively suggest that dormant tumor cells exhibit enhanced respiration and FAO.

### AMPK promotes mitochondrial biogenesis and FAO in dormant tumor cells

*In vitro*, MCF-7 and HCC-1428 cells underwent progressive metabolic alterations over time during hormone deprivation (HD) through treatment with medium containing 10% DCC-FBS (Fig. 3A). After 15 d of HD, ER+ BC cells had increased basal oxygen consumption rate (OCR), spare respiratory capacity, and ATP production rate (Fig. 3B-D). Additionally, 15-60 d of HD significantly increased mitochondrial membrane potential (Δ_Ψ_) and biogenesis in both cell lines (Fig. 3E/F). Consistent with these observations, HD increased FAO in ER+ BC cells (Fig. 3G).

**Fig. 3.**
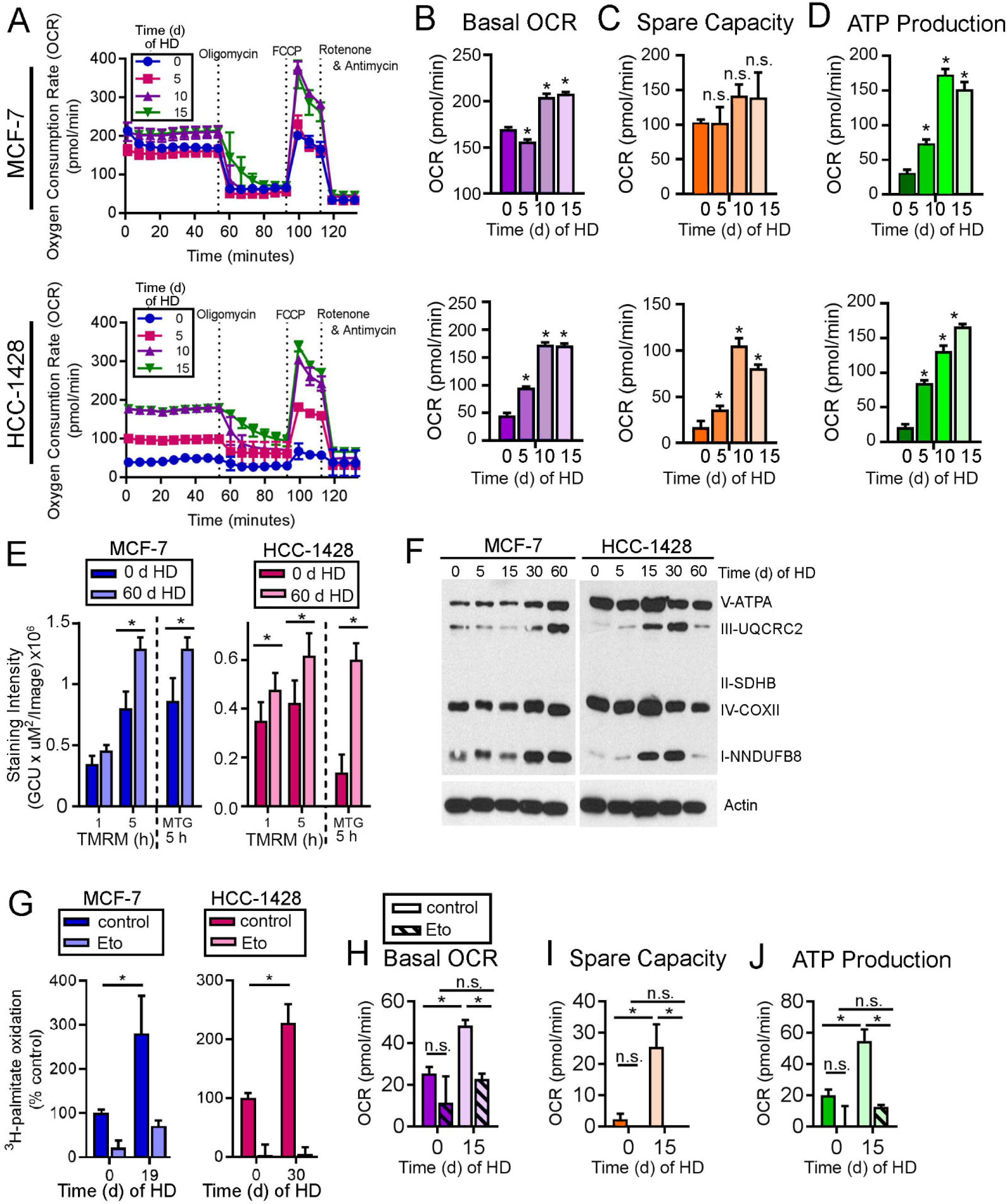
AMPK drives oxidative metabolism under HD conditions. **(A-D)** OCR was measured in cells treated with HD ±1 nM E2 for 15 d. Basal OCR (B), spare capacity (C), and ATP production (D) were calculated. **(E)** Cells were treated with HD ±1 nM E2, then stained with TMRM or MTG as indicators of mitochondrial membrane potential. **(F)** Lysates from cells treated with 0-60 d of HD were analyzed by immunoblot. **(G)** Cells were treated as in (A), then treated with medium containing ^3^H-palmitate ±20 μM etomoxir for 24 h prior to assay. FAO was normalized to HD + E2 control. **(H-J)** MCF-7 cells were treated with HD ±1 nM E2 for 15 d, ±200 μM etomoxir for 24 h prior to assay. OCR was measured as in (A-D). In (B-D) and (H-J), \**p*≤0.05 by Bonferroni multiple comparison-adjusted posthoc test vs. baseline unless indicated. In (E/G), * *t*-test *p*≤0.05. n.s.-not significant. In all quantitative experiments, data are shown as mean of triplicates +SD.

Since AMPKα2 is involved in oxidative metabolism and required for efficient FAO (21), we tested whether the HD-induced increases in respiration were driven by AMPK. Using siRNA against either AMPKα1 (*PRKAA1*) or AMPKα2 (*PRKAA2*) alone was insufficient to mitigate AMPK activity (data not shown). Dual knockdown of both AMPKα1/2 ablated AMPK activity (assessed by P-ACC loss) and reduced basal OCR and ATP production in ER+ BC cells under both HD and E2-stimulated conditions (Fig. S10), indicating the importance of AMPK in controlling oxidative metabolism.

Since dormant tumor cells exhibited transcriptional changes reflecting increased FAO (Fig. 2A), which is a major output of AMPK activity, we tested the role of FAO in HD-induced changes in oxidative metabolism. Treatment with etomoxir completely blocked HD-induced increases in basal OCR, spare capacity, and ATP production *in vitro* (Fig. 3H-J). These results suggest that, under HD conditions, AMPK activation drives metabolic adaptation to an increased oxidative state, primarily as a result of increased FAO.

### AMPK activity and fatty acid oxidation are essential for tumor cell survival following estrogen withdrawal

Since AMPK-directed mitochondrial respiration and FAO are increased following HD *in vitro*, we next tested the roles of these pathways in cancer cell persistence and growth. Genetic inhibition of AMPKα1/2 using siRNA significantly decreased MCF-7 cell number in HD conditions but not in the presence of E2 (Fig. 4A). Treatment with the AMPK-inhibiting compound dorsomorphin significantly decreased growth of 6/6 ER+ BC cell lines in HD and E2-stimulated conditions (Fig. 4B and data not shown).

**Fig. 4.**
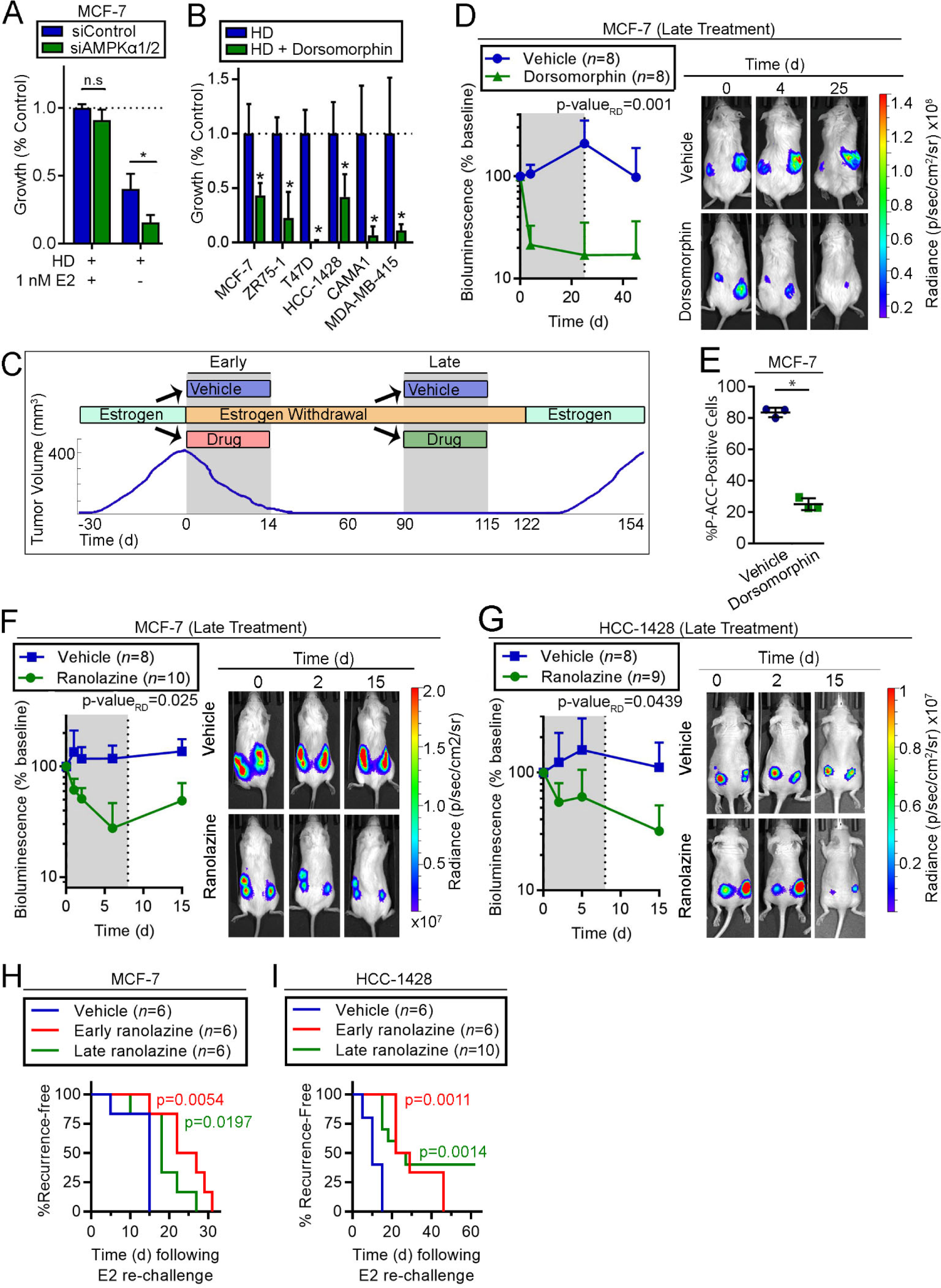
AMPK activity and fatty acid oxidation are essential for survival of dormant ER+ BC cells. **(A)** siRNA-transfected cells were treated with HD ±1 nM E2. Relative cell numbers measured after 7 d are shown as mean of triplicates +SD. **(B)** Cells were treated with HD ±5 μM dorsomorphin for 21-28 d and analyzed as in (A). **(C)** Schematic of experiments performed in (D-I). Ovx mice with E2-induced tumors were randomized to drug treatments either i) at the time of EW (“Early”), or ii) after 60-90 d of EW (“Late”). Tumor volume or bioluminescence was monitored before (Day 0), during (gray shading) and after drug treatments. **(D)** Mice with tumors that completely regressed after 90 d of EW were treated (“Late”) ± dorsomorphin for 25 d. Relative numbers of residual tumor cells were serially measured by bioluminescence imaging. Data are shown as mean +SD. Exposure-matched images from one representative mouse per group are shown. **(E)** Tumors from mice treated as in (D) were harvested 24 h after one dose. IHC was performed for P-ACC_Ser79_. Bars indicate mean of triplicates ± D. **(F/G)** Mice with tumors that completely regressed after 60 d of EW were treated (“Late”) ±ranolazine for 7 d, and analyzed as in (D). **(H/I)** Tumor-bearing mice were treated with EW +/-ranolazine: “Early” treatment was EW + ranolazine (14 d) administered to tumor-bearing mice, followed by 97 d of EW alone; after 90 d of EW, “Late” treatment was EW + ranolazine (7 d) administered to mice with completely regressed xenografts, followed by 7 d of EW alone. Mice without palpable tumors were then re-challenged with E2, and tumor recurrence was detected by palpation. Recurrence-free survival curves were compared to vehicle group by log-rank test. In (A/B/E), \**p≤*0.05 by *t*-test compared to control unless indicated. n.s.-not significant. In (D/F/G), bioluminescence signal values were compared by non-linear effect modeling.

In mice bearing dormant MCF-7/luciferase tumors (following 90 d of EW), “late” treatment with dorsomorphin rapidly decreased residual tumor cell number within 4 d (measured by bioluminescence imaging), suggesting induction of tumor cell death (Fig. 4C/D). Statistical comparison of rates of regression (*i.e.*, rate difference) of residual disease (*i.e.*, bioluminescence) between treatment groups showed that dorsomorphin had a significant effect compared to vehicle control (p-value_RD_=0.001). Following cessation of dorsomorphin treatment after 25 d, bioluminescence remained suppressed, indicating that A) the effects of dorsomorphin were durable, and B) luciferase expression/activity was not directly impeded by drug treatment. Dorsomorphin decreased P-ACC_Ser79_ IHC staining of tumor cells, confirming that AMPK activity was blocked (Fig. 4E). Since phosphorylation at Ser-79 inhibits ACC activity, which in turn promotes FAO (22), these data suggest that A) FAO was consequently inhibited by dorsomorphin, and B) AMPK activity, possibly by increasing FAO, is essential for survival of dormant ER+ breast tumor cells.

Ranolazine is FDA-approved for angina treatment and inhibits FAO. Thus, we tested whether ranolazine was effective for targeting of dormant ER+ breast tumor cells. “Late” treatment (as in Fig. 4C) of mice bearing dormant MCF-7/luciferase or HCC-1428/luciferase tumors (after 60 d of EW) with ranolazine for 7 d significantly increased the rate of decline of residual tumor burden measured by bioluminescence imaging (p-value_RD_=0.025 and 0.0439; Fig. 4F/G). These data suggest that dormant ER+ BC cells require FAO to survive. To test whether inhibition of FAO is effective when administered concurrently with EW, mice bearing E2-driven MCF-7 or HCC-1428 tumors were treated “early” (as in Fig. 4C) for 14 d with EW +/− one of three FAO inhibitors [ranolazine, etomoxir, or perhexiline, the latter two of which inhibit CPT1 (23–25)], followed by continued EW alone. All three FAO inhibitors hastened EW-induced tumor regression in one or both tumor models (Fig. S11) measured by both the rate of change in tumor volume (p-value_RD_) and the proportion of change in tumor volume with respect to baseline volume (*i.e.*, long-term treatment effect; p-value_LT_). We also tested whether “early” or “late” ranolazine treatment affects tumor re-growth potential, which indirectly reflects residual tumor cell burden. Mice pre-treated “early” with EW +/− 14 d of ranolazine that achieved complete tumor regressions (from Fig. S11A), and mice pre-treated with EW for 90 d followed by “late” ranolazine for 7 d, were re-challenged with E2. Both “early” and “late” ranolazine significantly delayed tumor recurrence compared to vehicle control (Fig. 4H/I). These data indicate that short-term FAO inhibition enhances the effects of EW on tumor regression and suppresses tumor regrowth.

### AMPK activation via metformin enhances the survival of ER+ residual disease during estrogen withdrawal

Since activation of FAO is a downstream effect of AMPK signaling, we sought to test the effects of pharmacologic AMPK activation on ER+ breast cancer cells. Metformin is an oral anti-hyperglycemic drug widely prescribed for the treatment of diabetes mellitus type 2. Although the mechanism by which metformin exerts anti-diabetic effects remains controversial and is likely multi-faceted, it is generally accepted that a main cellular consequence of metformin is activation of AMPK to modulate metabolic and lipid signaling (26). Based primarily on A) epidemiologic evidence suggesting that metformin has anti-cancer properties in diabetic patients (27), and B) preclinical data showing that metformin slows cancer cell and tumor growth (28, 29), metformin is being heavily tested clinically for the treatment of a variety of cancer types in non-diabetic patients. Reports have thus far yielded conflicting results (30–33), which may be due in part to the variable pharmacokinetic properties and therapeutic effects of metformin in humans (34). Conflictingly, data also support a tumor-promoting role for AMPK. P-ACC levels in primary tumors are associated with shorter overall survival in patients with non-small cell lung cancer and head and neck squamous cell carcinomas (35, 36). AMPK activation is associated with disease recurrence in prostate cancer, the mechanism of which was linked to PGC-1α signaling and subsequent mitochondrial biogenesis and function (37). Our data indicated that AMPK-directed metabolism drives survival of residual ER+ breast tumor cells during EW (Fig. 4). Thus, we tested whether AMPK activation via metformin affects growth of ER+ BC cells and tumors during HD and EW, respectively.

Little is known about the concentrations of metformin that reach cancerous tissues in humans. *In vitro* evidence identifying metformin as an effective anti-cancer agent was generated with supra-pharmacologic concentrations of metformin (5-50 mM); achievable plasma concentrations in humans are in the low micromolar range. Thus, we first performed a pharmacokinetic study in mice to obtain an understanding of achievable tissue concentrations of metformin. Mice were administered ∼5 mg/d metformin via drinking water for 6 wk. Resultant steady-state plasma concentrations of metformin were 4.27 ± 1.19 μM (mean ± SD; Fig. S12A), which are in agreement with the preclinical findings of Dowling *et al*. and reflect human steady-state plasma concentrations of 0.08-6.5 μM (38). This dose in mice elicited growth-inhibitory effects on MCF-7 xenografts (p-value_RD_<0.001) and activated AMPK in tumor tissues (Fig. S12B/C). Metformin concentrations in tissues (tumor, liver, muscle) range from 0.47 to 1.14 μM (Fig. S12A). In parallel, our studies in cultured MCF-7 cells revealed that treatment with 1 mM metformin- the lowest dose that maximally induced AMPK activation-provided intracellular concentrations of 49.1 ± 10.8 μM (Fig. S12D/E). Treatment with 1 mM metformin significantly inhibited growth of ER+ BC cells under E2-stimulated conditions (Fig. S12F); this dose was selected for further *in vitro* studies.

*In vitro*, treatment with 1 mM metformin significantly rescued growth of 6/6 ER+ BC cell lines in HD conditions for ≥3 wk (Fig. 5A). The ability of metformin to rescue such growth was ablated upon siRNA-induced knockdown of AMPKα1/2, suggesting that metformin rescues growth in HD conditions via AMPK activation (Fig. 5B).

**Fig. 5.**
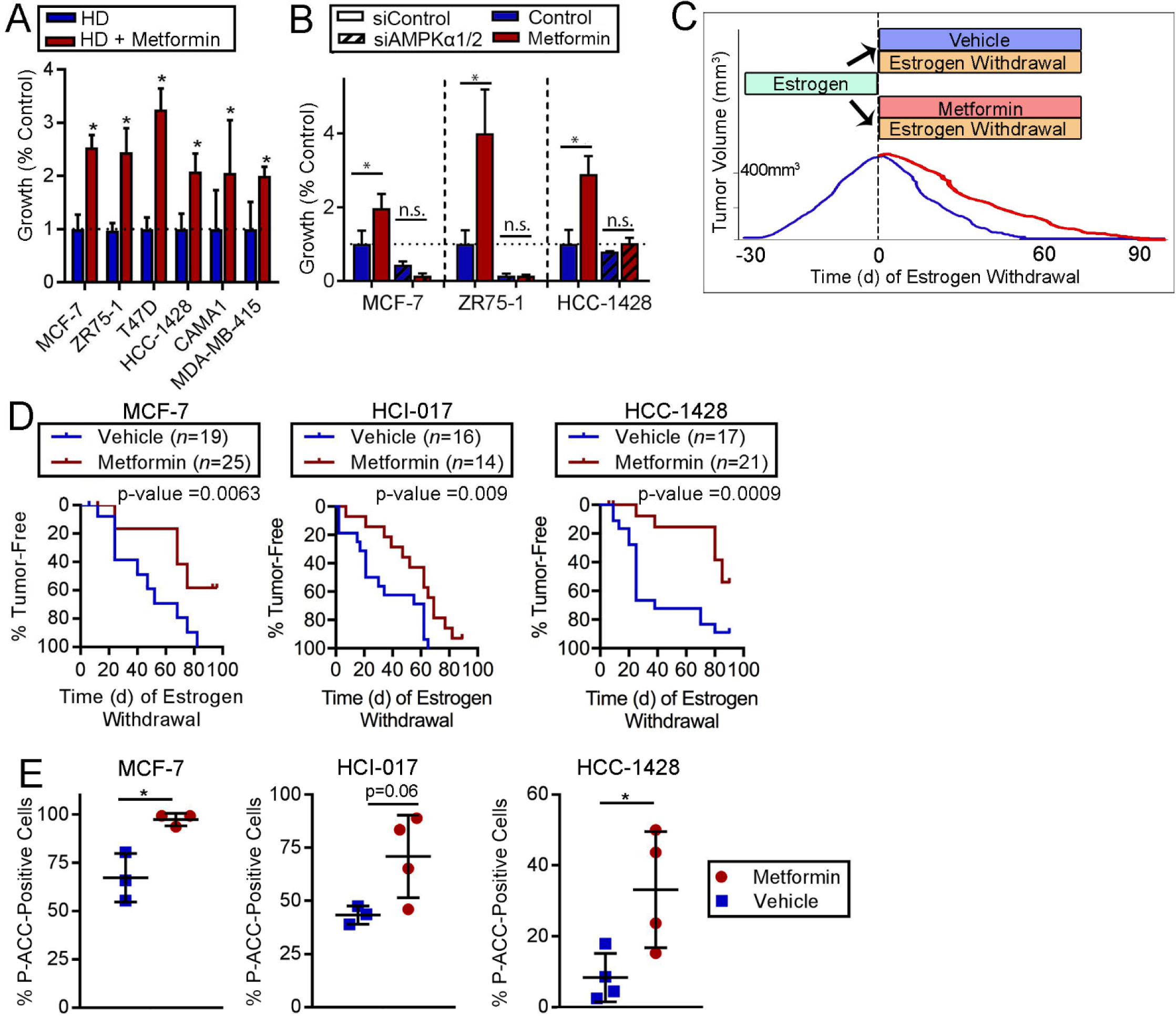
AMPK activation via metformin promotes survival of ER+ BC cells following estrogen withdrawal. **(A)** Cells were treated with HD ±1 mM metformin. Relative viable cell numbers after 21-28 d were compared to control by *t*-test (\**p≤*0.05). **(B)** siRNA-transfected cells were treated for 7 d then analyzed as in (A). \**p*≤0.05 by Bonferroni multiple comparison-adjusted posthoc test. n.s.-not significant. In (A/B), data are presented as mean of triplicates +SD. **(C)** Schematic of experiment performed in (D). Ovx mice with E2-driven tumors (400 mm^3^) were treated with EW ± metformin (∼5 mg/d via drinking water) for 90 d. Tumor volumes were monitored by caliper measurements. **(D)** Proportions of tumors that completely regressed over time were compared by log-rank test. **(E)** Tumors harvested from mice after 12 d (MCF-7) or 6 d (HCI-017, HCC-1428) of treatment with EW ± metformin were analyzed by IHC for P-ACC_Ser79_. \**t*-test *p≤*0.05.

To test the effects of metformin on ER+ breast tumor cell persistence during EW, we treated mice bearing E2-driven tumors with EW ± metformin and monitored tumor volume (Fig. 5C); this is analogous to the clinical scenario of treatment with AI therapy ± metformin. In MCF-7, HCI-017, HCC-1428, and MDA-MB-415 models, concurrent treatment with metformin slowed EW-induced tumor regression compared to EW/placebo controls (all log-rank p<0.05; Figs. 5D and S13-14). Metformin did not significantly alter levels of serum insulin, serum free fatty acids, or blood glucose (Fig. S15). Increased P-ACC staining was observed in tumors obtained from mice treated with metformin for 6-12 d (Figs. 5E and S16). These data indicate that metformin enhances ER+ breast tumor cell persistence and AMPK activation during EW.

Interestingly, metformin did not rescue HCC-1500 tumors from regression induced by EW *in vivo*, or HCC-1500 cells from HD *in vitro* (Fig. S17A/B). HCC-1500 cells have relatively high levels of AMPKα2 protein and AMPK activity compared to MDA-MB-415, HCC-1428, and MCF-7 cells (Fig. S17C). Furthermore, metformin did not alter P-ACC levels in HCC-1500 cells *in vitro* or EW-treated tumors *in vivo* (Fig. S17D/E); these observations explain in part why metformin did not enhance cell persistence upon HD/EW in this model, and emphasize the dependence upon AMPK activation for metformin effects in ER+ BC cells and tumors.

### Fatty acid oxidation sustains ER+ breast cancer cells during estrogen deprivation

In E2-stimulated conditions, 24 h of treatment with metformin enhanced markers of glucose and fatty acid metabolism, cell cycle inhibition, and autophagy. In HD conditions over a time-course of 15 d, metformin enhanced expression of FAO enzymes and mitochondrial respiratory proteins (Fig. 6A). Treatment with the FAO inhibitor etomoxir ablated the ability of metformin to sustain growth of HCC-1428 and MCF-7 cells in HD conditions (Fig. 6B), suggesting that metformin drives FAO to promote hormone-independent growth/persistence.

**Fig. 6.**
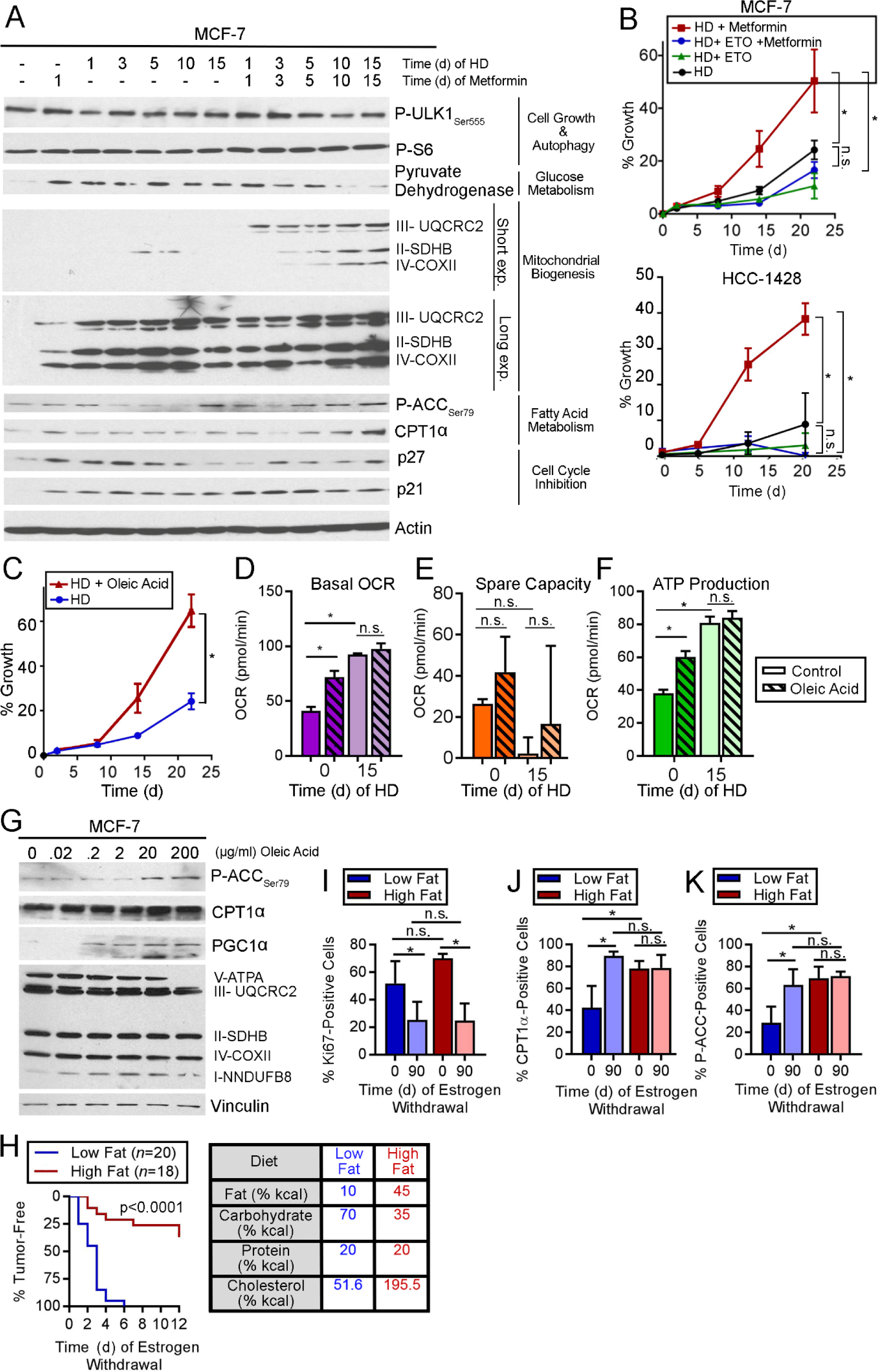
Dietary fat promotes survival of ER+ BC cells following estrogen withdrawal. **(A)** Immunoblot analysis of lysates from cells treated with HD ±1 nM E2 and metformin. **(B/C)** Cells were treated with HD ±200 μM etomoxir or 1 mM metformin (B), or ±300 μg/mL oleic acid (C). Relative viable cells were serially measured. **(D-F)** OCR was measured from cells treated with HD ±1 nM E2 for 15 d, then treated ±200 μg/mL oleic acid for 5 h. Basal OCR (D), spare capacity (E), and ATP production (F) were measured. **(G)** Lysates of cells treated ± oleic acid for 24 h were analyzed by immunoblot. **(H)** Ovx mice pre-conditioned to either high-fat or low-fat diet for 2 wk were injected orthotopically with MCF-7 cells and supplemented with E2 (with diet continuation). When tumors reached ∼400 mm^3^, E2 was withdrawn. Tumor volumes were serially measured. Data are presented as proportions of mice reaching complete tumor regression over time. Groups were compared by log-rank test. Caloric sources in diets are noted at right. **(I-K)** IHC for Ki67, CPT1α, or P-ACC_Ser79_ using tumors harvested after 0 or 90 d of EW. In (B/C), \**p*≤0.05 by repeated measures ANOVA. In (D-F) and (I-J), \**p*≤0.05 by Tukey’s multiple comparison-adjusted posthoc test vs. baseline unless otherwise indicated. n.s.- not significant. In (B-F) and (I-J), data are shown as mean of triplicates ± SD.

Since FAO sustains ER+ BC cells during HD *in vitro*, and adipocytes can acts as an energy source for tumor cells through provision of fatty acids (39), we tested whether exogenous fatty acid elicits similar effects. Despite enhancing MCF-7 cell growth in HD conditions (Fig. 6C), treatment with free oleic fatty acid did not alter basal OCR, spare capacity, or ATP production after 15 d of HD, possibly because HD cells had saturated FAO or respiratory activity (Fig. 6D-F). Interestingly, oleic acid treatment increased basal OCR, ATP production, and levels of P-ACC and PGC-1α in E2-stimulated conditions (Fig. 6D-G), suggesting that cancer cells can respond to increased fatty acid abundance (*i.e.*, fuel) by increasing FAO (*i.e.*, catabolism).

In mice bearing E2-driven MCF-7 tumors, pre-conditioning with a high-fat diet slowed EW-induced tumor regression (p-value_RD_<0.001; Fig. S18), as was observed with metformin (Figs. 5D and S13-14). While pre-conditioning with a low-fat diet provided EW-induced complete tumor regression in 100% (20/20) of cases within 6 wk, a high-fat diet allowed complete regression in only 38% (7/19) of cases within 12 wk (log-rank p<0.0001; Fig. 6H). IHC analysis of tumor specimens suggested that differences in tumor response to EW were not due to dietary pre-conditioning effects on tumor cell proliferation (Ki67), but rather due to high-fat diet-induced increases in tumor cell FAO (*i.e.*, increased CPT1α and P-ACC) at the start of EW on Day 0 (Figs. 6I-K and S19).

## Discussion

Herein, we described the development of preclinical models of dormancy in ER+ BC. In an EW state, as would occur in patients treated with AI therapy, small proportions of ER+ breast tumor cells persisted in a growth-suppressed state for extended periods of time, and retained tumor-forming capacity. Such dormant tumors exhibited features of both population-level and cellular dormancy. Molecular analyses revealed upregulation of AMPK levels and activity in dormant tumors compared to E2-driven or acutely EW tumors. Dormant tumor cells were dependent upon AMPK activity for persistence that was enhanced by metformin, and AMPK inhibition reduced dormant tumor cell burden. In culture, HD stimulated a metabolic switch to an increased oxidative state with dependence upon FAO; AMPK activity drove these metabolic alterations. Similarly, dormant tumor cells exhibited markers of increased respiration and dependency upon FAO, whereupon FAO inhibitors reduced dormant tumor cell burden as well as hastened EW-induced tumor regression.

The biology of dormant tumor cells that can cause cancer recurrence is not well-understood due in part to the lack of understanding of the location(s) and context(s) of such cells in patients. DTCs have been found in bone marrow and CTCs have been found in blood because these tissues can be sampled in patients; whether occult cancer cells exist in other tissues/organs remains an open question. As an initial investigation into mechanisms driving the persistence of dormant tumor cells, we evaluated orthotopic xenograft models of human ER+ BC in immunodeficient mice; potential translational limitations of these models are the mammary adipose tissue microenvironment, and the microenvironment of regressed tumors. In humans, ∼30% of ER+/HER2− BC recurrences observed within 10 yr of follow-up occur locally in the ipsilateral breast, while the majority of recurrences occur in distant metastatic sites (1). Determining whether similar mechanisms of dormancy and therapeutic sensitivity occur in cancer cells in mammary adipose tissue vs. distant sites of metastasis (e.g., bone, liver, lung, brain) will require the generation of organ-specific models of metastatic ER+ BC. We established small E2-driven tumors (∼9 mm in diameter) and induced regression by EW to study persistent dormant cancer cells; this paradigm reflects the clinic scenario of occult residual/micrometastatic disease in patients treated with AI therapy that is below the limit of detection (∼7 mm) of PET/CT (40). The extracellular microenvironment of regressed tumors may differ from that of isolated DTCs, which could also affect dormancy signalling and drug sensitivity.

How dormant tumor cells interact with and respond to a functional (human) immune system may provide clues to improve therapy; however, the model systems used herein are immunodeficient. Emerging evidence suggests that transcriptional programs that control dormancy (*i.e.*, growth suppression, angiogenic switch) might also coordinate an immune-evasive phenotype (41), offering a potential therapeutic strategy to sensitize dormant cells to immunotherapies. Work by Malladi *et al*. showed that latent metastatic human cancer cells implanted in nude mice were maintained in a slow-cycling dormant state by autocrine WNT inhibition, which caused downregulation of signals required for natural killer cell-mediated clearance (42). Future studies involving animal models with humanized immune systems would aid in investigating the roles of immune evasion and surveillance in cancer dormancy.

Precision medicine is based upon the premise that using the right treatment, for the right patient, at the right time will maximize therapeutic index. However, ‘time’ (*i.e.*, stage during progression of a disease) is not always considered during drug development. History provides examples of FDA-approved drugs that showed efficacy against (clinically evident) recurrent/metastatic solid tumors, but which failed to increase recurrence-free survival when tested in adjuvant settings (in patients with no evidence of residual disease) (43–49). Such disparate findings between adjuvant and recurrent/metastatic settings may be partly attributable to biological differences: the biology and microenvironments (*e.g.*, vasculature, stroma, extracellular matrix, inflammation, hypoxia) of established tumors likely differ substantially from those of dormant tumor cells; thus, drug sensitivity may also differ. Drug treatments are generally most beneficial when tumor burden is low [*i.e*., adjuvant setting following surgical resection of measurable tumor(s)], so the typical drug development paradigm has been to demonstrate clinical efficacy against overt recurrent/metastatic disease, then clinically test the drug in the adjuvant setting; the gap in this paradigm is preclinical testing of the drug in an “adjuvant setting.” Herein, we present a preclinical platform to test therapeutic strategies that target dormant residual disease in the background of EW therapy for ER+ BC. It is hoped that such preclinical testing would become standard practice prior to initiation of a costly adjuvant clinical trial that will take years to yield results, particularly with ER+ BC due to long durations of dormancy and times to recurrence.

Based on preclinical molecular analyses of dormant vs. growing ER+ breast tumors, we identified AMPK activation as a driver of dormant cancer cell persistence, which led to evaluation of metformin. Preclinical studies have shown that metformin elicits anti-proliferative effects in a variety of cancer cell lines *in vitro* and *in vivo*, which is thought to occur partly by suppression of mTORC1 signaling (26,28,29). Conversely, metformin slowed EW-induced regression of ER+ breast tumors and promoted persistence/growth of ER+ BC cells during HD (Figs. 5A/D and S13-14), suggesting that AMPK activation promotes ER+ BC cell survival during the stress of EW. In line with these findings, AMPK has been shown to promote cancer cell survival during stress induced by nutrient starvation (16, 50). Our collective findings highlight ‘timing’ as an important consideration in precision medicine and therapeutic development. Extrapolating these findings to the human condition, metformin use concurrent with anti-estrogen therapy may promote the survival of residual endocrine-sensitive ER+ BC cells, which could increase risk of recurrence. Thus, the timing of metformin therapy in patients with ER+ BC may impact clinical benefit, and timing needs to be considered when interpreting clinical trial results. Although AMPK inhibition reduced ER+ breast tumor cell burden under EW conditions (Fig. 4A-D), this may be a difficult therapeutic strategy to develop clinically because an AMPK inhibitor could also promote tumor growth. Focusing on the metabolic alterations induced by AMPK activation, we found that dormant ER+ breast tumor cells were dependent upon FAO, which may be a more practical therapeutic target. Indeed, FAO inhibitors are clinically approved for angina treatment and could be repurposed as cancer therapeutics.

## Acknowledgements

We thank Alana Welm for sharing the HCI-017 tumor model, Jonas Dehairs and Ali Talebi for guidance on assaying FAO, and Craig Tomlinson for advice on animal diets. We thank the following Norris Cotton Cancer Center Shared Resources for assistance: Mouse Modeling, Clinical Pharmacology, Pathology, Genomics & Molecular Biology, and Immunoassays & Flow Cytometry. Funding support was provided by the Mary Kay Foundation (036-15 to TWM), the Foundation for Women’s Wellness (Research Grant to TWM), Friends of the Norris Cotton Cancer Center (Scholar Award to TWM), NIH (R01CA211869 to TWM, R01CA200994 to TWM, R01CA126618 to WBK, F31CA220936 to RAH, F30CA216966 to KS, and Dartmouth College Norris Cotton Cancer Center Support Grant P30CA023108), and the Burroughs Wellcome Big Data in the Life Sciences Training Program (fellowship to JDW). RNA-seq data are available at the NCBI SRA website under accession number PRJNA491455.

