## Supplementary material for "AMPK activation by metformin promotes survival of dormant ER+ breast cancer cells": Suppl Figures S1-S19 and Methods

**Supplemental information for Hampsch *et al*.**

**This file is organized as:**


A. **Legend for Supplemental Table S1 (page 1).**
B. **Supplemental Figures S1-S19 with legends (pages 2-20).**
C. **Supplemental Methods (pages 22-29).**
D. **Supplemental References Cited (pages 29-30).**


### A. Legend for Supplemental Table.

**Table S1- Normalized mRNA counts for NanoString PanCancer Pathways Analysis.** RNA was isolated from FFPE MCF-7 tumor specimens from mice treated with EW for 0, 3, or 82 d. Hybridization and subsequent sample analysis were performed per manufacturer's protocol. Raw mRNA counts were normalized using nSolver software, and normalized counts are presented below.

**B. Supplemental Figures S1-S19 with legends.**

**Figure S1. ER+ breast cancer cells lie dormant during estrogen depletion *in vivo* and retain tumor-initiating capacity.** MCF-7 cells were s.c. injected into ovx mice, and mice were immediately randomized to s.c. implantation of an E2 pellet (black) or sham control (blue/red). After 10 wk, control mice remaining were then randomized to E2 pellet (blue) or control (red). Tumor volumes were measured twice weekly.

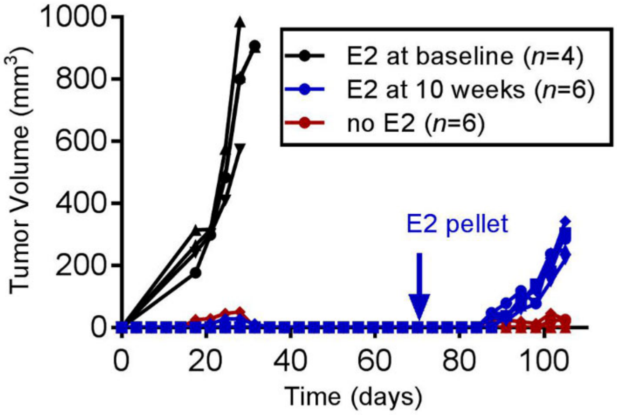

**Figure S2. Individual tumor regression and growth curves for mice with estrogen-withdrawn and estrogen-retreated xenografts. (A-E)** Ovx mice bearing bilateral E2-driven orthotopic tumors measuring ~400 mm$^3$ were treated with estrogen withdrawal starting on Day 0. Tumor volumes were measured twice weekly. Each line represents one tumor. Summarized data are shown in Fig. 1B. **(F-H)** Ovx mice bearing E2-driven orthotopic tumors were treated with estrogen withdrawal as above. After 90 d of estrogen withdrawal, mice were retreated with E2 via s.c. pellet on Day 0. Tumor volumes were measured twice weekly. Each line represents one tumor.

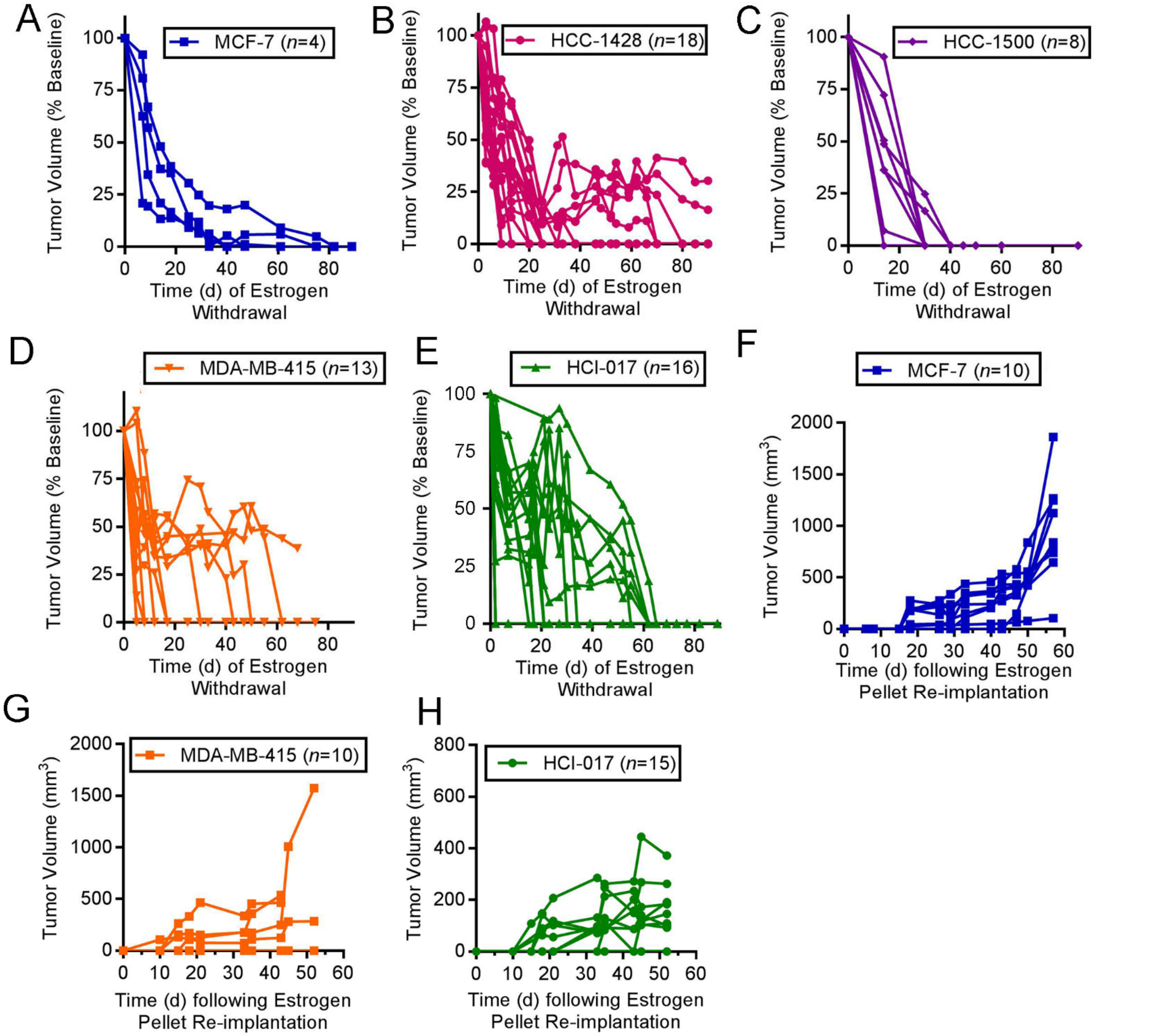

**Figure S3. Immunohistochemical analysis of Ki67, cleaved caspase 3/7, and ER in ER+ breast cancer xenografts.** Representative images for Ki67 **(A)**, cleaved caspase 3/7 **(B)**, and ER **(C)** IHC staining of FFPE xenografts are shown. Tumor specimens were harvested from ovx mice following 0, 6, or 90 d of estrogen withdrawal. Summarized data for (A) and (B) are shown in Fig. 1F-H. Summarized data for (C) are shown in (D). Data are shown as mean of triplicates + SD. *p≤0.05 by t-test. n.s.- not significant.

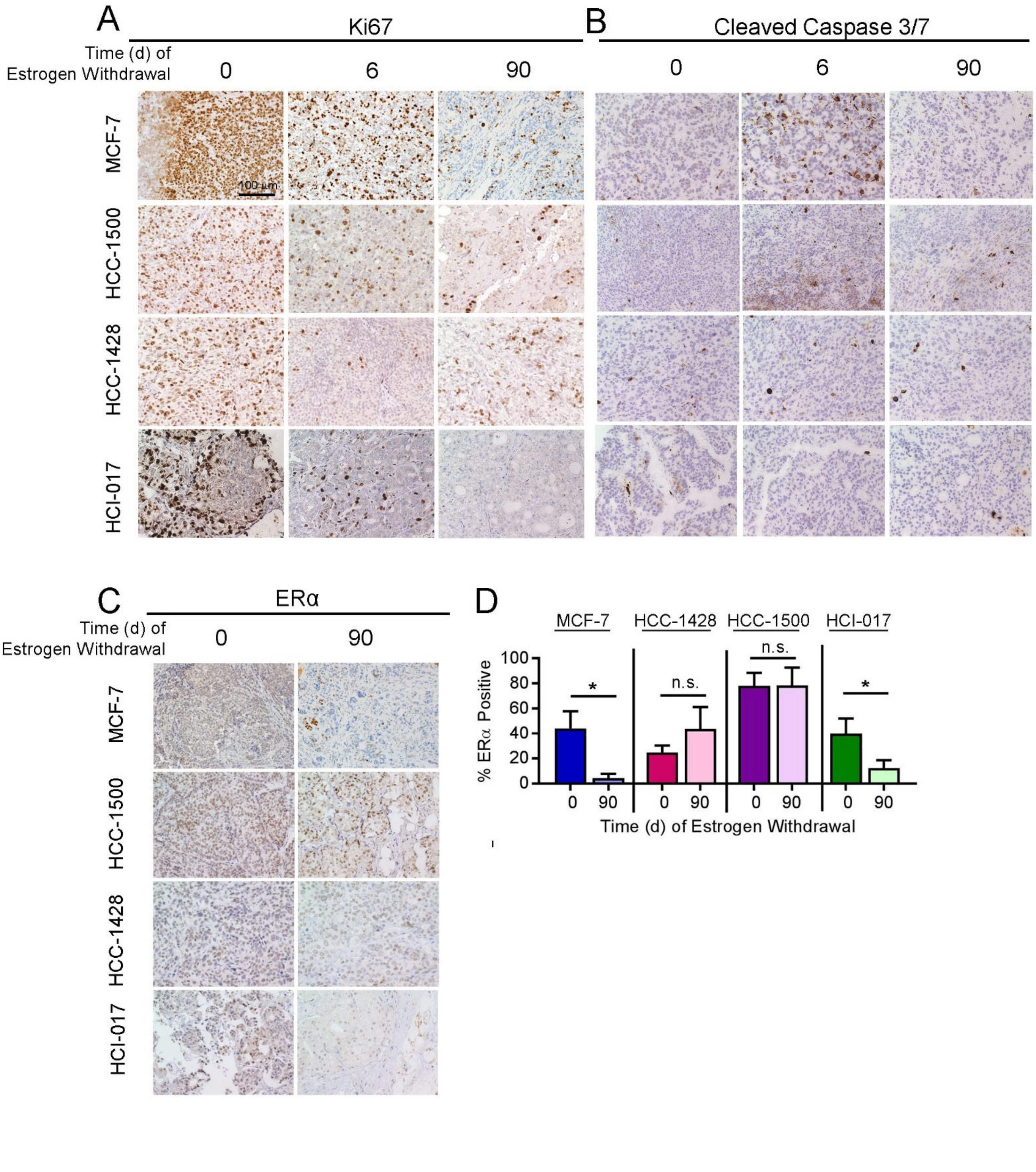

**Figure S4. Gene Set Variation Analysis (GSVA) of RNA sequencing data from ER+ breast cancer xenografts.** RNA extracted from MCF-7 or HCC-1428 tumors after 0 or 90 d of estrogen withdrawal (*n*=3-4 tumors/time point) was analyzed by sequencing. Gene set variation analysis (GSVA) of whole-transcriptome expression profiles of clinically dormant tumors (Day 90) compared to baseline (Day 0) was performed using unsupervised sample-wise enrichment analysis of common metabolic and signaling gene sets [selected from the Hallmarks (HM), Gene Ontology (GO), Reactome (RM), or Motif Gene Sets (C3) collections]. Vertical dotted line indicates adjusted p=0.2 threshold, which is below the recommended cutoff of p=0.25 for significance in Gene Set Enrichment Analysis (1). Summarized data are shown in Fig. 2A.

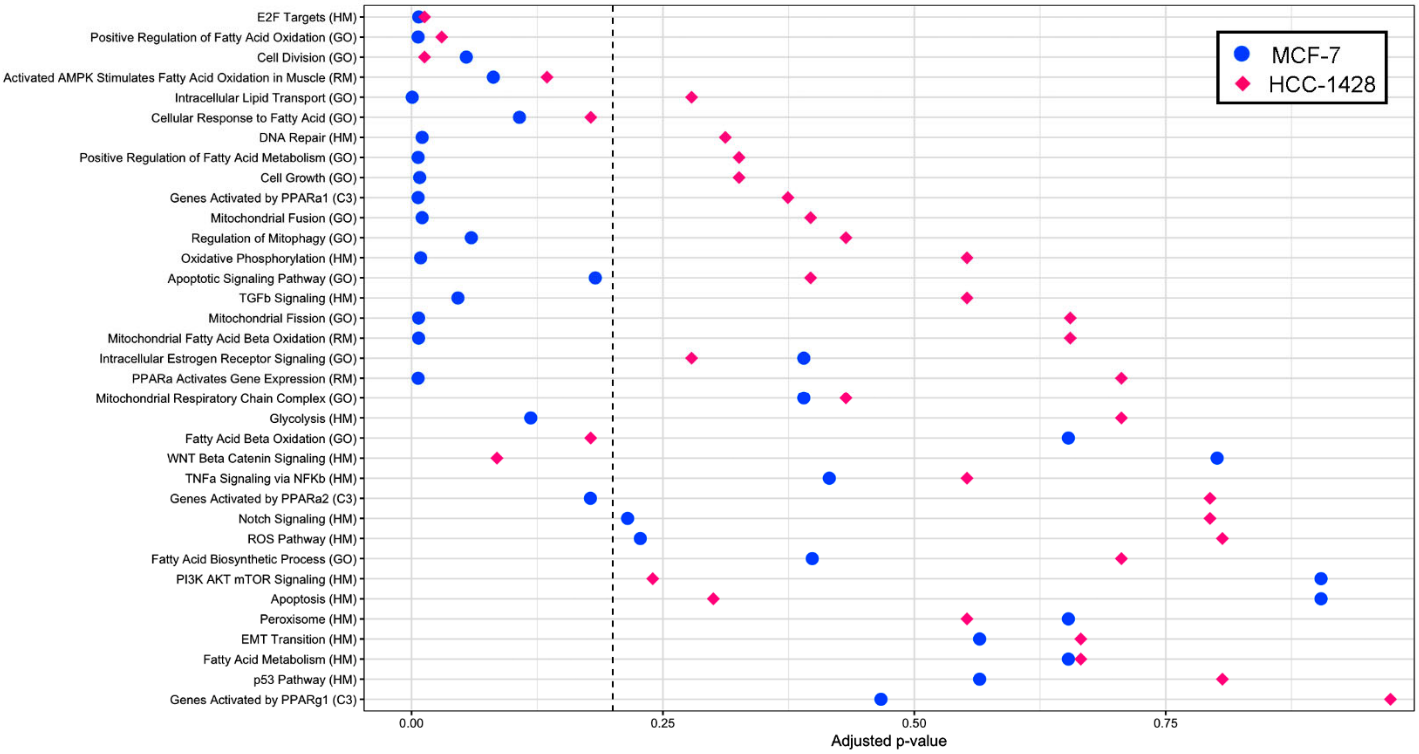

**Figure S5. RNA sequencing analysis of ER+ BC xenografts.** RNA was extracted from macrodissected sections of FFPE MCF-7 **(A)** or HCC-1428 **(B)** tumors harvested from ovx mice after 0, 6, or 90 d of estrogen withdrawal (EW; *n*=3-4 tumors/time point), and analyzed by sequencing. Genes that were significantly differentially expressed ($q \leq 0.05$, $|log_2FC|>1.0$) between Day 6 (acute EW) and Day 0 (baseline) (blue and yellow circles), or between Day 90 (dormant) vs. Day 0 (baseline) (red and green circles) were identified; numbers of these genes are indicated in Venn diagrams. *AMPKα2 (PRKAA2) mRNA was upregulated at Day 90, but not Day 6, compared to baseline in both tumor models. GSVA results are shown in Figure 2A and Figure S4.

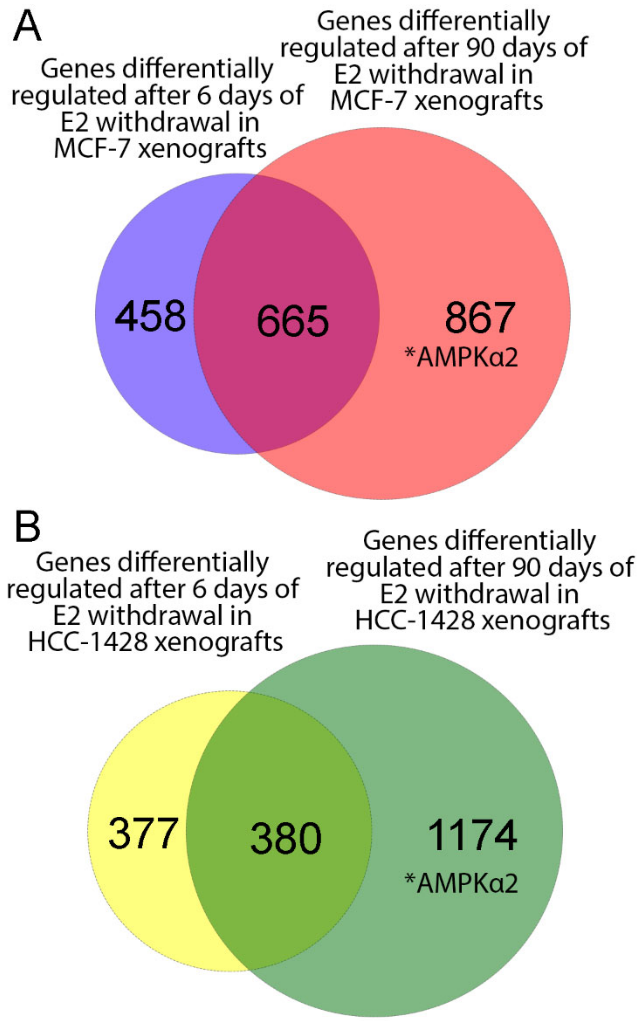

**Figure S6. Immunohistochemical analysis of AMPKα2 and P-ACC in ER+ breast cancer xenografts.** Representative images for AMPKα2 **(A)** and P-$ACC_{Ser79}$ **(B)** IHC staining of FFPE xenografts are shown. Tumor specimens were harvested from ovx mice following 0, 6, or 90 d of estrogen withdrawal. Summarized data are shown in Fig. 2C/D.

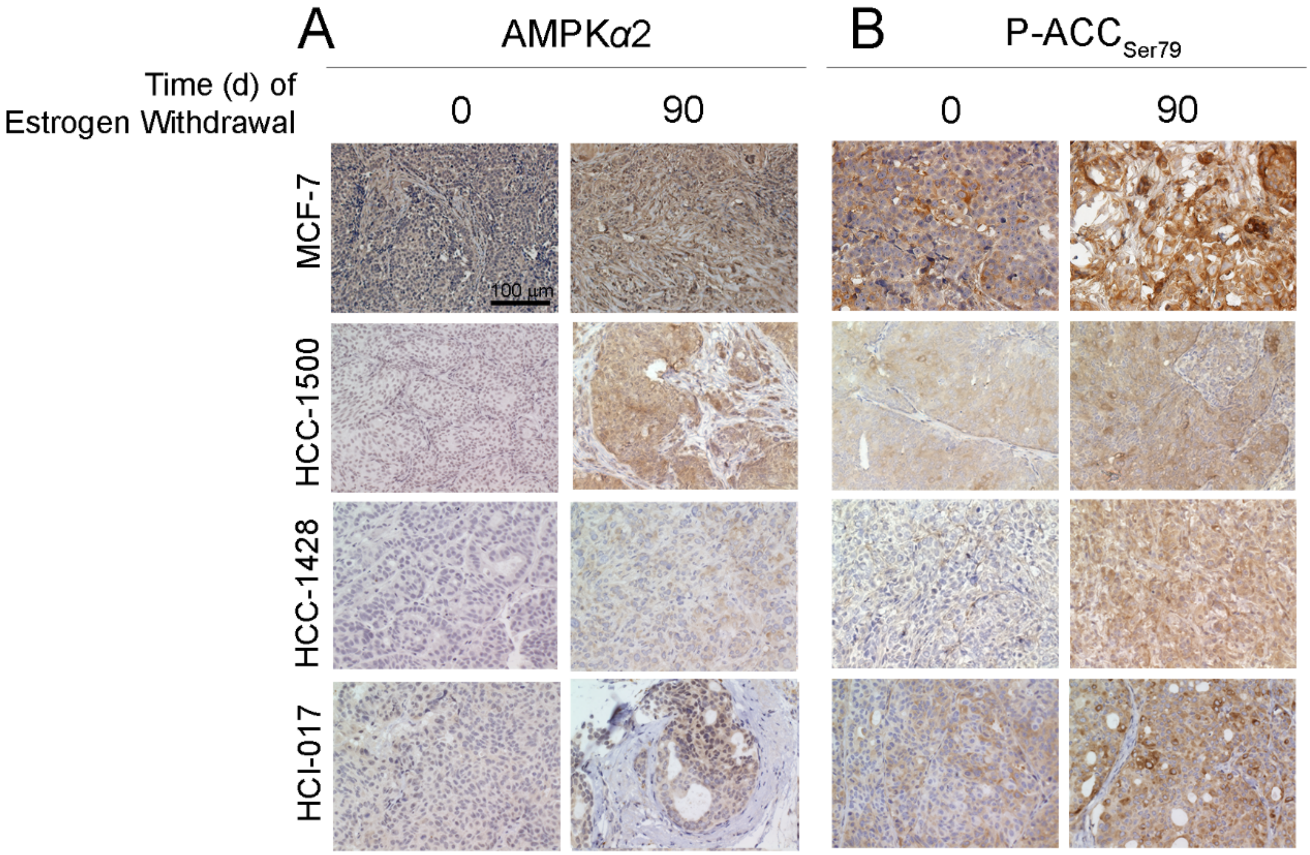

**Figure S7. Effects of dormancy on mTORC1 activation in ER+ breast tumors. (A)** MCF-7 tumors were harvested from ovx mice after 0 or 90 d of estrogen withdrawal (EW). FFPE tumors specimens were analyzed by IHC for phospho-S6. Representative images are shown. Proportions of positively-stained cells were measured using HaloVelocity software in 3 images per tumor; values were averaged within each tumor (3 tumors per group). Quantification is shown on right. Bars indicate mean ± SD. Groups were compared using *t*-test. **(B)** In a separate experiment, GFP+ MCF-7 tumors were harvested from ovx mice after 0, 3, 6, 12, or 90 d of EW (3 tumors per time point). Tumors were enzymatically digested into single-cell suspensions, stained using phospho-S6 antibody, and analyzed by flow cytometry. Proportions of GFP+ cells that were P-S6-positive were measured. Data are shown as mean of triplicates ± SD. *$p \leq 0.05$ by Bonferroni multiple comparison-adjusted posthoc test.

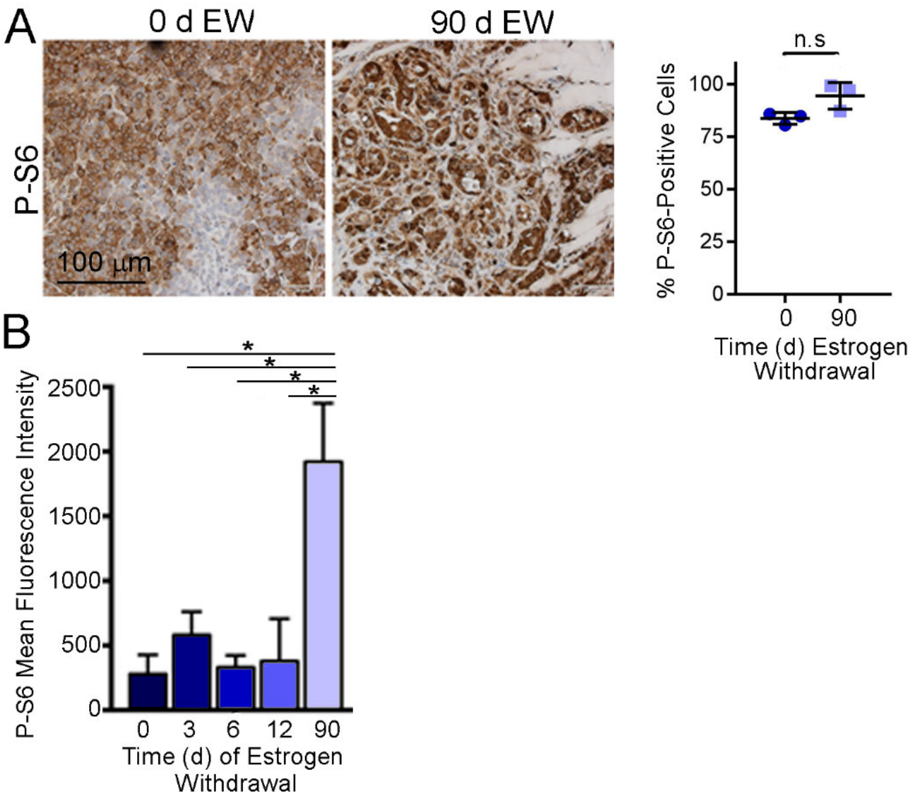

**Figure S8. Residual ER+ breast tumor cells exhibit a transcriptional signature of FAO following presurgical anti-estrogen treatment.** Human ER+ breast tumor etomoxir *t*-statistics were calculated using the 915 genes non-cell cycle-related gene signature of etomoxir response from MCF-7 cells. There were 863 (GSE20181), 629 (GSE71791), and 632 (GSE111563) non-cell cycle-related genes available on each platform. Groups were statistically analyzed by repeated measures ANOVA followed by Tukey's multiple comparison-adjusted posthoc test between time points (A) or paired *t*-test (B/C). Red bars indicate mean values.

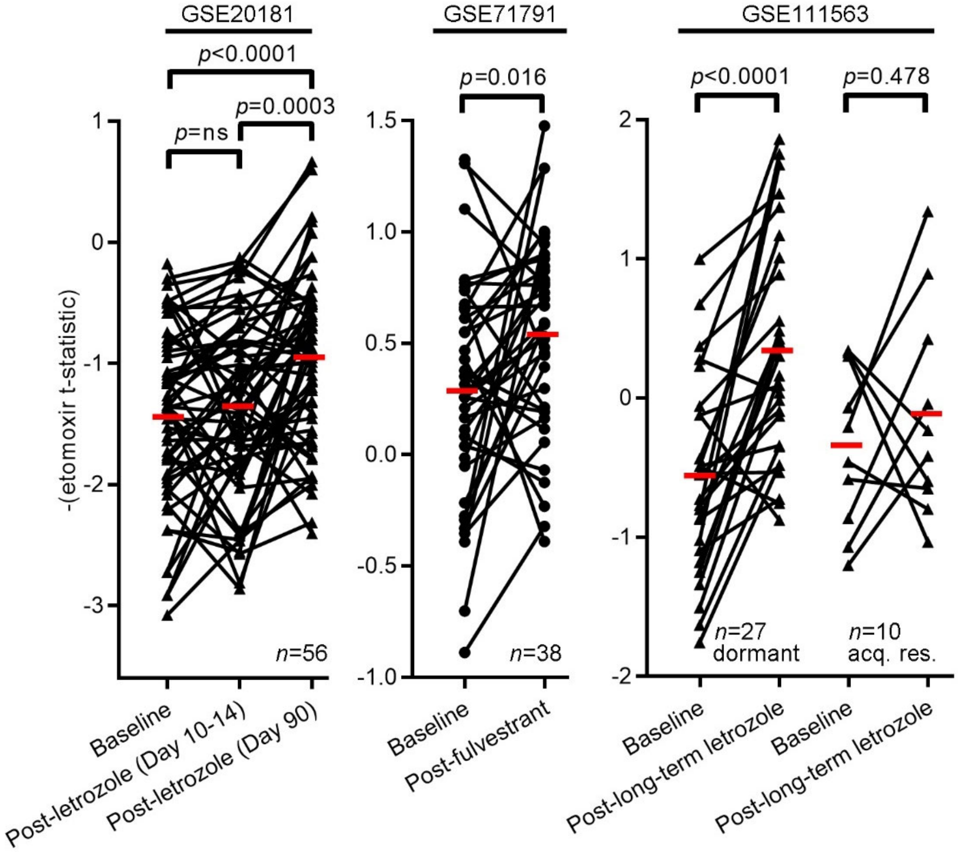

**Figure S9. Clinically dormant tumor cells exhibit markers of increased mitochondrial content and activity. (A)** Normalized mRNA read counts for mitochondria-encoded genes extracted from RNA-seq data. **(B-C)** Mitochondria in tumor specimens harvested after 0 or 90 d of EW were stained with TOM20-AF594 by immunofluorescence. Mitochondrial count (B) and length (C) per cell (from ≥30 cells) were calculated in each of 3 tumors per group. *$p \leq 0.05$ by *t*-test. (D) Representative IHC images for CPT1α staining of tumors. Summarized data are shown in Fig. 2H.

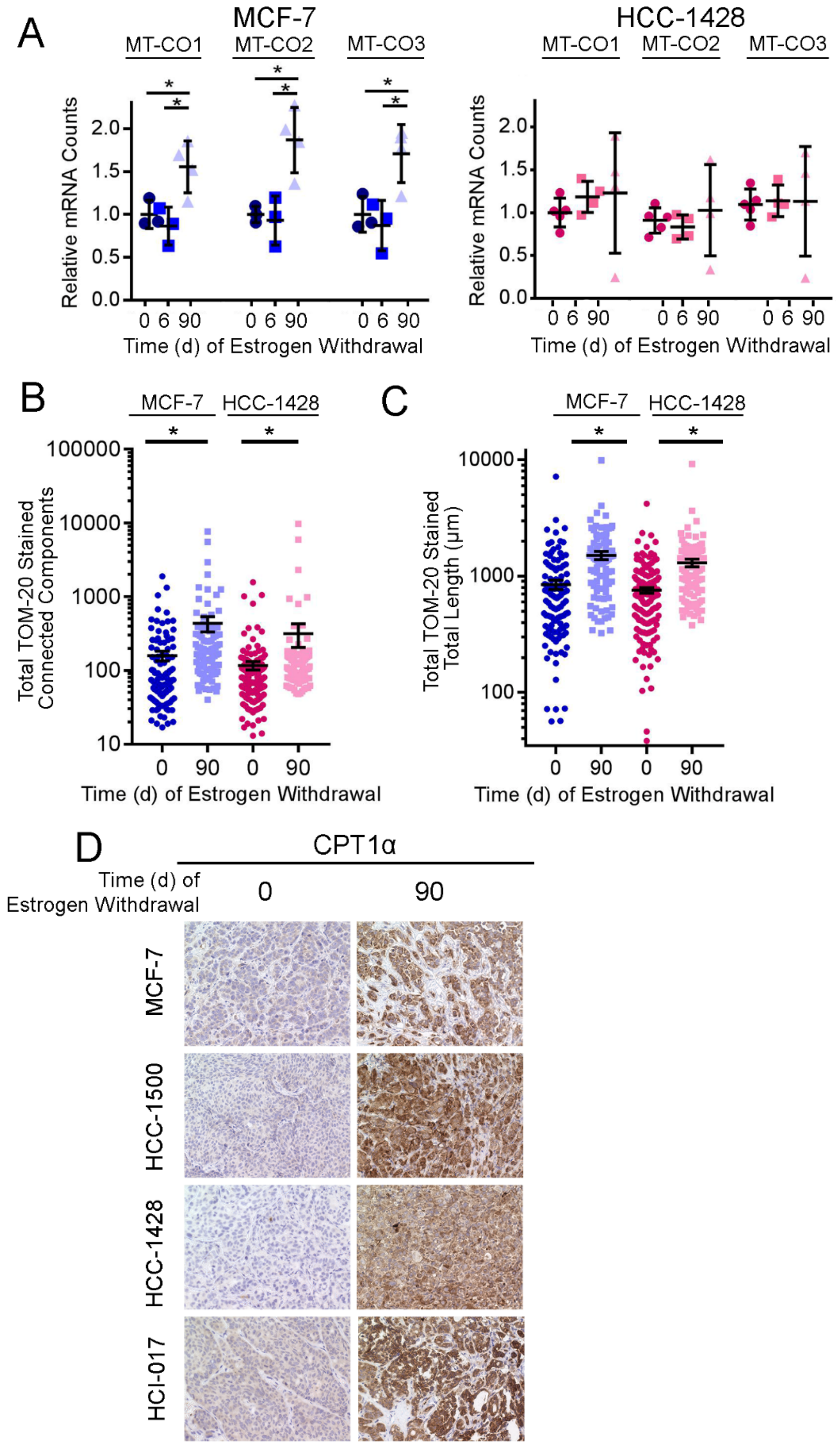

**Figure S10. AMPK is required for hormone deprivation-induced increases in cellular respiration in ER+ breast cancer cells. (A)** Cells were transiently transfected with siRNA targeting AMPKα1 and/or AMPKα2, or non-silencing control. Three days later, lysates were harvested and analyzed by immunoblot. **(B-D)** MCF-7 cells were pre-treated with hormone-depleted (HD) medium ± 1 nM E2 for 12 d, then transfected with siRNA against AMPKα1 and AMPKα2, or non-silencing control. Cells were maintained in growth medium or HD medium for an additional 3 d, then assayed for cellular respiration. Basal OCR (B), spare capacity (C), and ATP production (D) measurements are shown as mean of triplicates + SD. **p*≤0.05 by Bonferroni multiple comparison-adjusted posthoc test. n.s.- not significant.

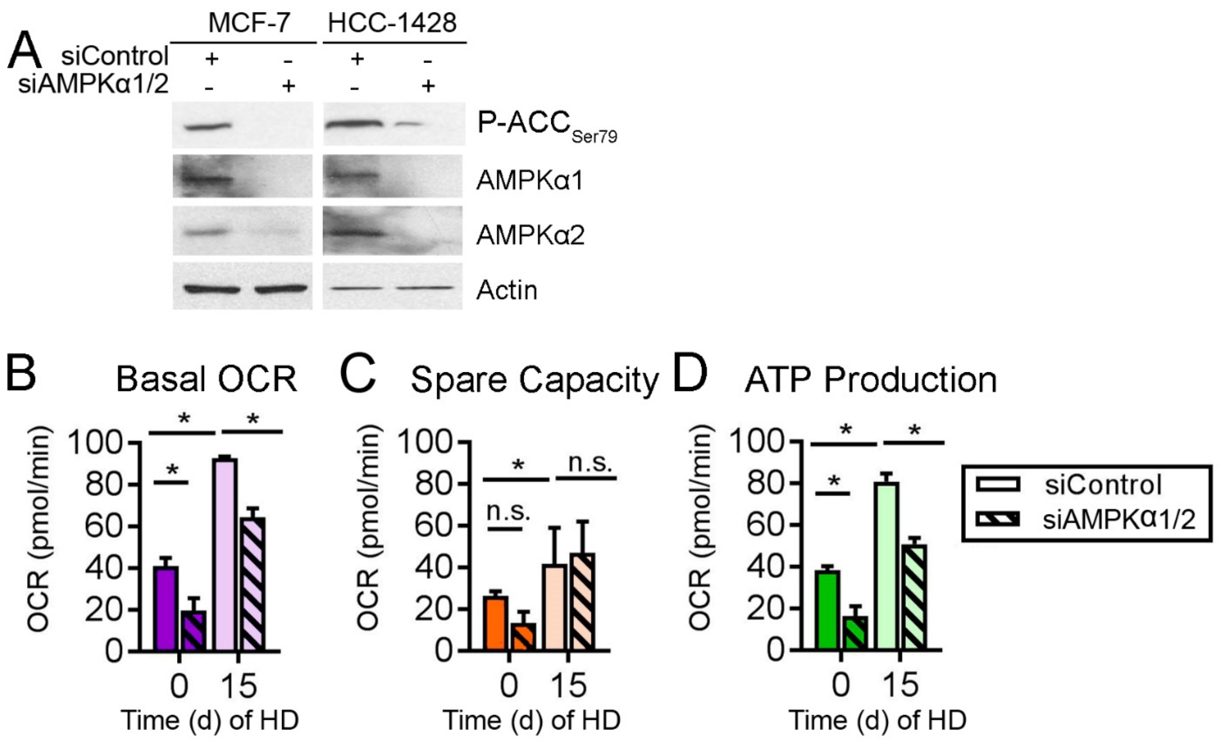

**Figure S11. Inhibition of fatty acid oxidation hastens regression of ER+ breast tumors following estrogen withdrawal.** Ovx mice bearing orthotopic bilateral E2-driven breast tumors (~400 $mm^3$) were randomized to treatment with estrogen withdrawal (EW) plus 14 d of drug treatment with ranolazine (50 mg/kg QD i.p.), etomoxir (50 mg/kg QD i.p.), perhexiline (15 md/kg BID i.p.), or vehicle control ("Early Treatment" as shown in Fig. 4C). Tumor volumes were measured twice weekly. Gray shading indicates the 14-day drug treatment period. EW continued after drug treatment ceased. Tumor volumes are shown as mean + SD. One drug treatment vs. vehicle is shown in each plot for visualization; vehicle group is the same in all plots within each model [(A) shows MCF-7; (B) shows HCC-1428]. Groups were compared using non-linear effect modeling depending on regression patterns. $\text{p-value}_{RD}$ reflects differences between rates of tumor regression; $\text{p-value}_{LT}$ reflects differences in tumor volume in the long term. For the MCF-7 model, tumors quickly regressed below the threshold for measurement, so long-term treatment effects could not be statistically evaluated.

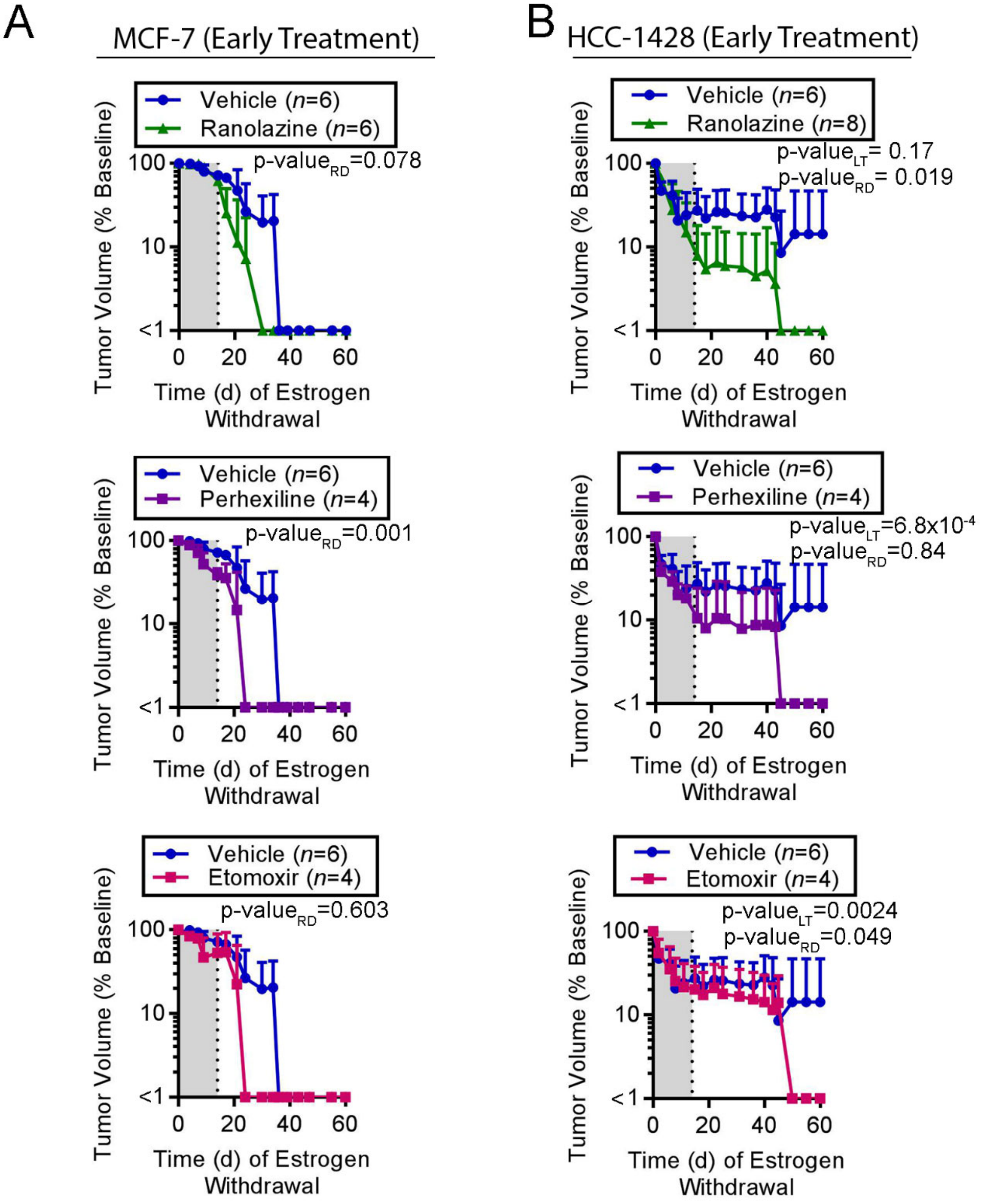

**Figure S12. Pharmacologic analysis of metformin in ER+ breast cancer cells and tumors in mice. (A)** Mice (*n*=5) were treated with metformin (~5 mg/d via drinking water) for 6 wk. Mice were euthanized. Blood was harvested and used to extract plasma. Tumor, muscle, and liver tissues were harvested. Plasma and tissues specimens were analyzed by LC-MS/MS to measure metformin concentrations. Bars indicate mean ± SD. **(B)** Ovx mice bearing E2-driven MCF-7 tumors were treated with EW for ~90 d, which caused complete regression of most tumors. In mice without palpable (residual) tumors, E2 supplementation was re-applied, and mice were simultaneously randomized to treatment ± metformin (~5 mg/d) on Day 0. Tumor volumes were measured twice weekly. Groups were compared using non-linear effect modeling depending on regression patterns. p-value$_{RD}$ reflects differences between rates of tumor growth. **(C)** Lysates from tumors harvested at the study endpoint in (B) were analyzed by immunoblot. **(D)** Cultured MCF-7 cells were treated with a dose range of metformin in growth medium for 24 h. Lysates were collected and analyzed by immunoblot. P-ACC and P-ULK1 were used as readouts of AMPK activity. **(E)** Cells were treated as in (D), then collected for measurement of concentrations of metformin (reported as mean ± SD of biological triplicates). **(F)** Cells were treated with hormone-depleted medium (HD) + 1 nM E2 ± 1 mM metformin for 21-28 d. Cells were then stained with crystal violet, and staining intensity was quantified. Data are shown as mean of triplicates + SD. *$p \leq 0.05$ by *t*-test compared to each control.

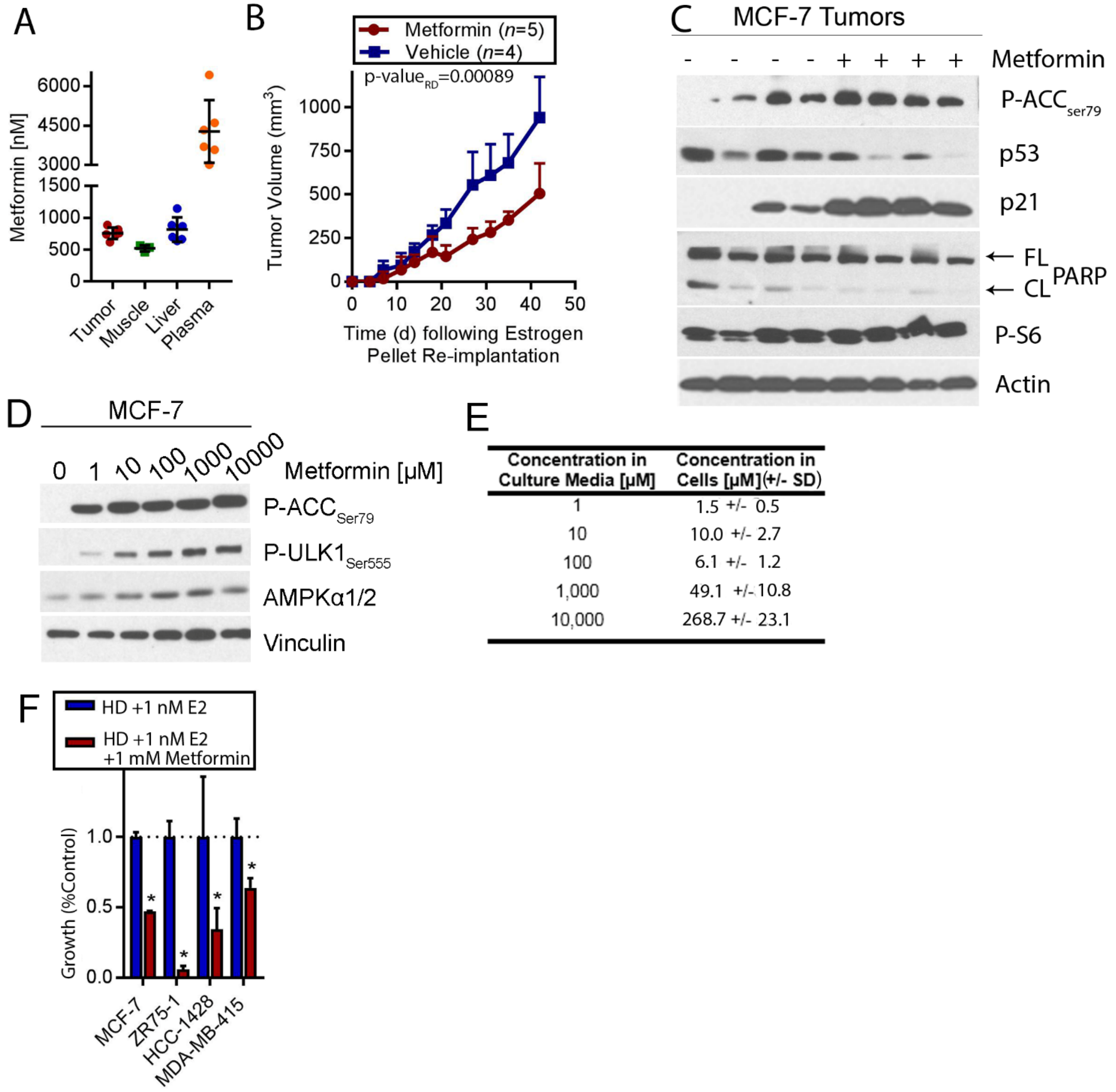

**Figure S13. Metformin prevents estrogen withdrawal-induced regression of MDA-MB-415 ER+ breast tumors. (A)** Ovx mice were orthotopically injected bilaterally with MDA-MB-415/luciferase cells, and implanted s.c. with an E2 pellet. Relative tumor cell numbers were serially measured by bioluminescence imaging. Tumors became palpable, and E2 pellets were removed on Day 117. Most tumors then regressed to a non-palpable state (indicated by gray shading). Each line represents an individual tumor. A representative bioluminescence image is shown at right after 90 d of EW. **(B)** Ovx mice bearing E2-driven MDA-MB-415/luciferase tumors (~400 mm$^3$) were treated with EW, and simultaneously randomized to treatment ± metformin (~5 mg/d via drinking water). Proportions of tumors that completely regressed (*i.e.*, non-palpable) over time are indicated. Kaplan-Meier curves were compared by log-rank test.

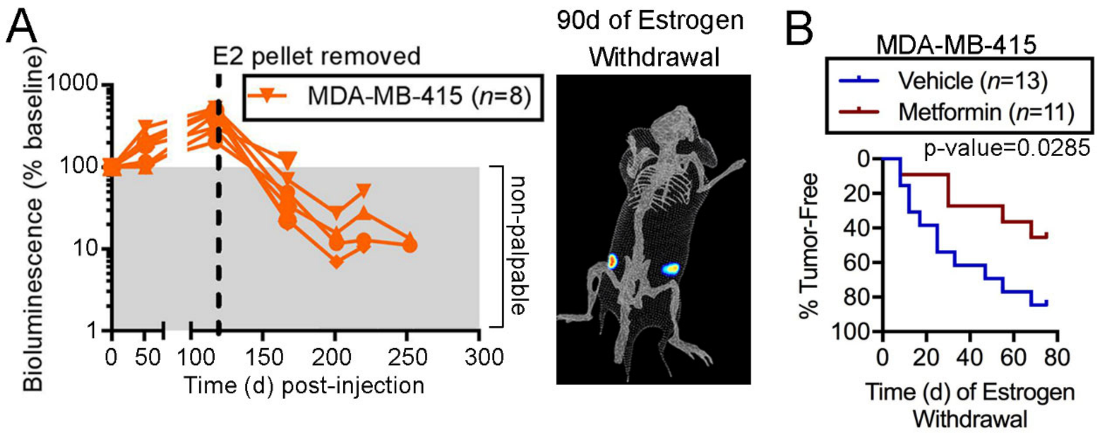

**Figure S14. Metformin prevents estrogen withdrawal-induced regression of ER+ breast tumors.** Ovx mice bearing orthotopic bilateral E2-driven tumors (~400 $mm^3$) were treated with EW, and simultaneously randomized to treatment ± metformin (~5 mg/d via drinking water). Tumor volumes are shown as mean ± SD. Groups were compared using non-linear effect modeling: p-value$_{RD}$ reflects differences between rates of tumor growth/regression; p-value$_{LT}$ reflects differences in tumor volume (*i.e.*, long-term treatment effect, or the proportion of tumor volume following treatment with respect to baseline tumor volume). Since metformin treatment was continued for the duration of the experiment, we focused on long-term treatment effects. However, MCF-7 and HCI-017 vehicle-treated tumors quickly regressed to below the limit of detection; thus, we could not measure long-term treatment effects (to generate p-value$_{LT}$), and instead measured short-term treatment effects to assess differences in rates of regression (to generate p-value$_{RD}$).

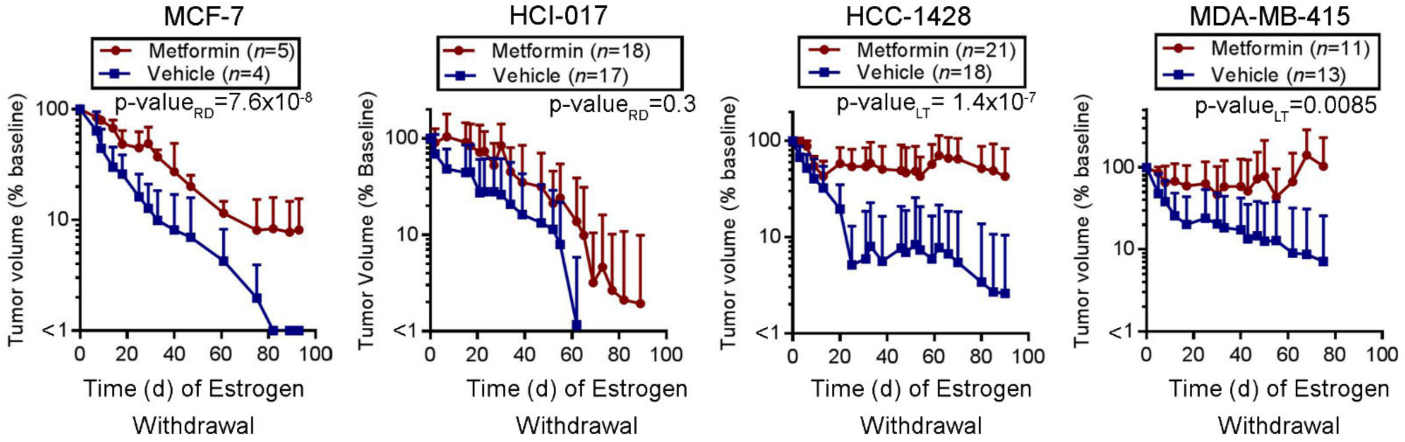

**Figure S15. Metformin treatment does not significantly alter fasting levels of blood glucose, serum insulin, or serum free fatty acids in mice.** Mice were treated ± metformin (~5 mg/d via drinking water) for 6 wk. Mice were fasted for 12 h overnight, then bled retroorbitally. Blood was used to measure concentrations of glucose **(A)**, or processed to extract serum. Serum was used to measure concentrations of insulin **(B)** and free fatty acids **(C)**. Each point represents one mouse. Bars indicate mean ± SD. Groups were compared by *t*-test. n.s.- not significant.

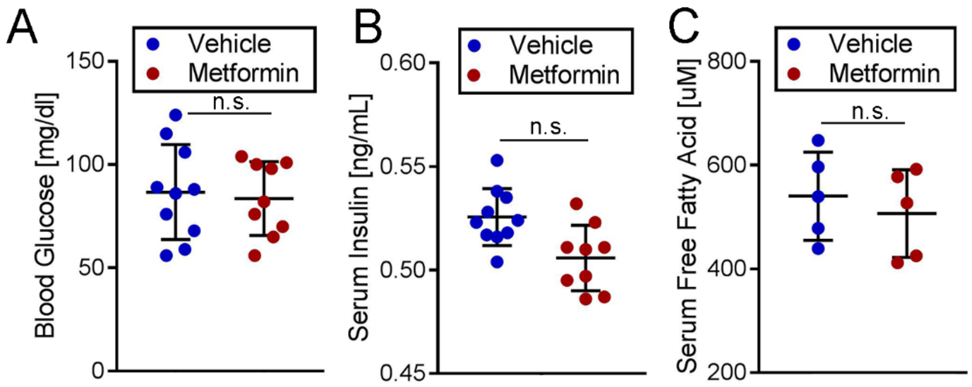

**Figure S16. Immunohistochemical analysis of P-ACC in ER+ breast tumors from mice treated ± metformin.** Representative images for P-ACC$_{Ser79}$ IHC staining of FFPE xenografts are shown. Tumor specimens were harvested from ovx mice following 12 d (MCF-7) or 6 d (HCC-1500, HCC-1428, HCI-017) of EW. For the duration of those 6-12 d, mice were randomized to receive metformin or vehicle control. Summarized data are shown in Fig. 5E.

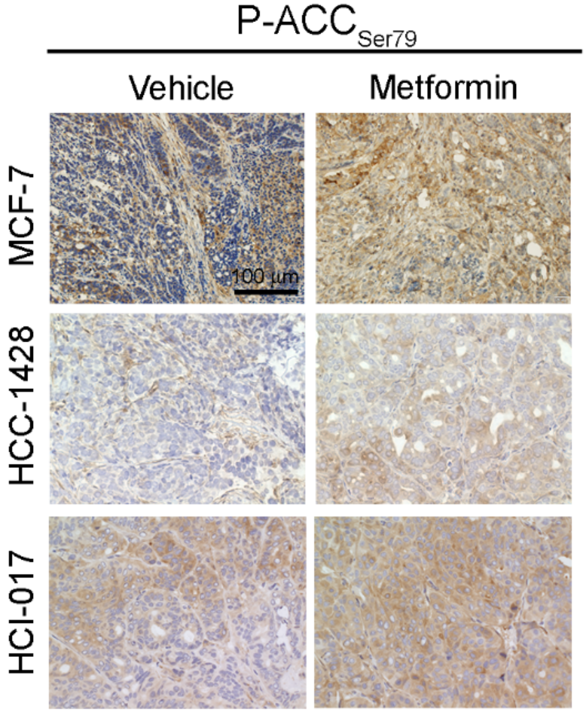

**Figure S17. AMPK modulation does not alter growth/survival of HCC-1500 ER+ breast cancer cells or tumors. (A)** HCC-1500 cells were treated with hormone-depleted medium (HD) ± 1 mM metformin for 21-28 d in triplicate. Relative cell numbers were quantified by Incucyte. Data are shown as mean + SD. **(B)** Ovx mice bearing orthotopic bilateral E2-driven HCC-1500/luciferase tumors (~400 $mm^3$) were treated with EW, and simultaneously randomized to treatment ± metformin (~5 mg/d via drinking water). Proportions of mice with complete tumor regression (*i.e.*, non-palpable) are indicated. Kaplan-Meier curves were compared by log-rank test. **(C)** Lysates from ER+ breast cancer cell lines were analyzed by immunoblot. P-$ACC_{Ser79}$ and P-$ULK1_{Ser555}$ were used as readouts of AMPK activity. **(D)** Cells were treated ± 1 mM metformin, then lysates were analyzed by immunoblot. **(E)** Ovx mice bearing orthotopic bilateral E2-driven HCC-1500 tumors (~400 $mm^3$) were treated with EW ± metformin (~5 mg/d) for 6 d. Tumors were harvested, FFPE, and sections were stained by IHC for P-$ACC_{Ser79}$. Proportions of positively-stained cells were measured in 3 microscopic fields (200x magnification) per tumor, and the mean value was used for each tumor (shown as one point in graph). Horizontal bars indicate mean ± SD. In (A/E), groups were compared by *t*-test. *$p \leq 0.05$. n.s.- not significant.

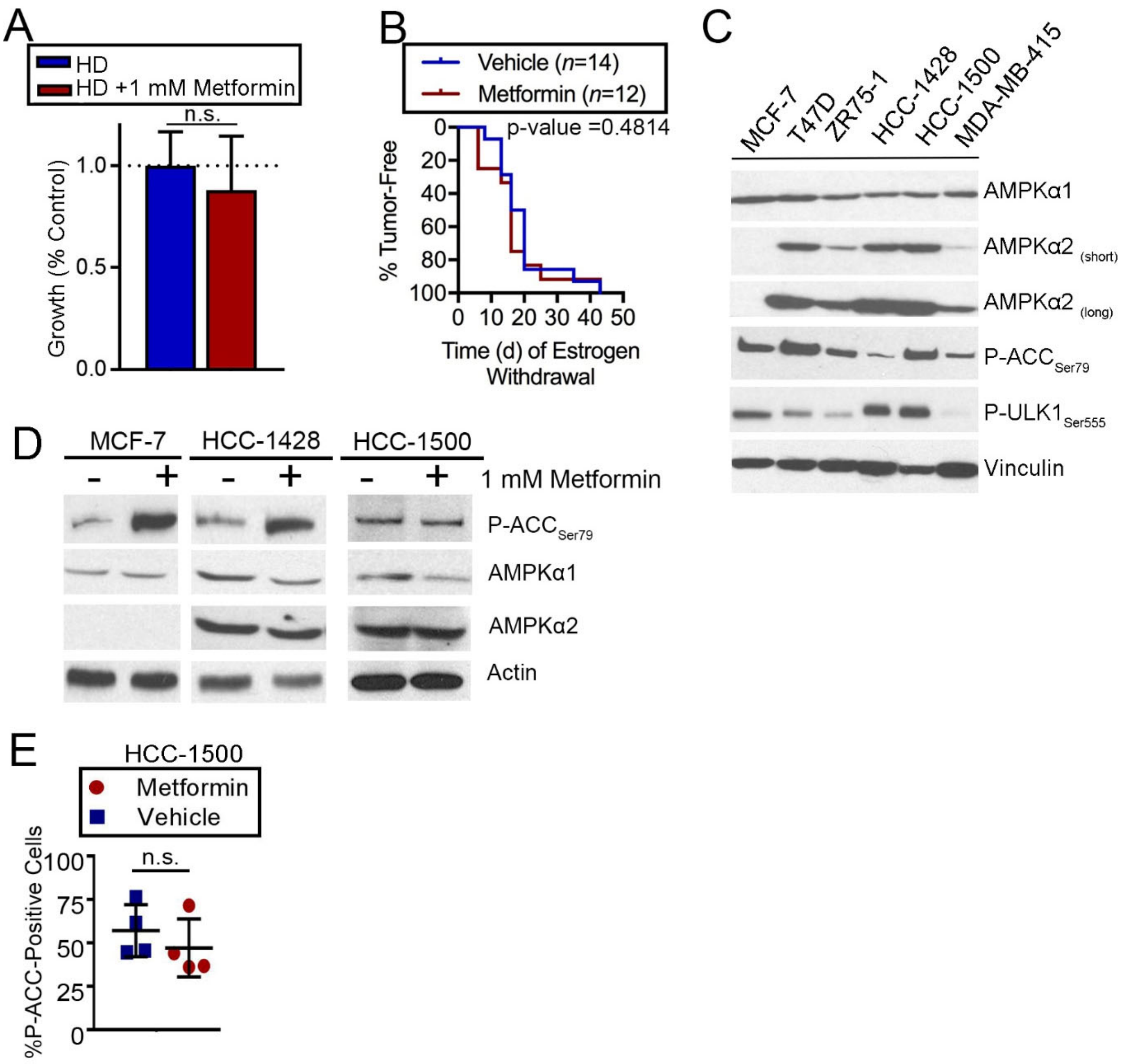

**Figure S18. High-fat diet prevents estrogen withdrawal-induced regression of ER+ breast tumors.** Ovx mice were conditioned to either a high-fat or low-fat diet for 2 wk, then injected orthotopically bilaterally with MCF-7 cells and implanted s.c. with an E2 pellet (with diet continuation). When tumors reached ~400 $mm^3$, mice were treated with EW (with diet continuation). Tumor volumes were serially measured, and groups were compared using non-linear effect modeling. p-value$_{RD}$ reflects differences between rates of tumor regression. Growth/regression curves of individual tumors are shown below.

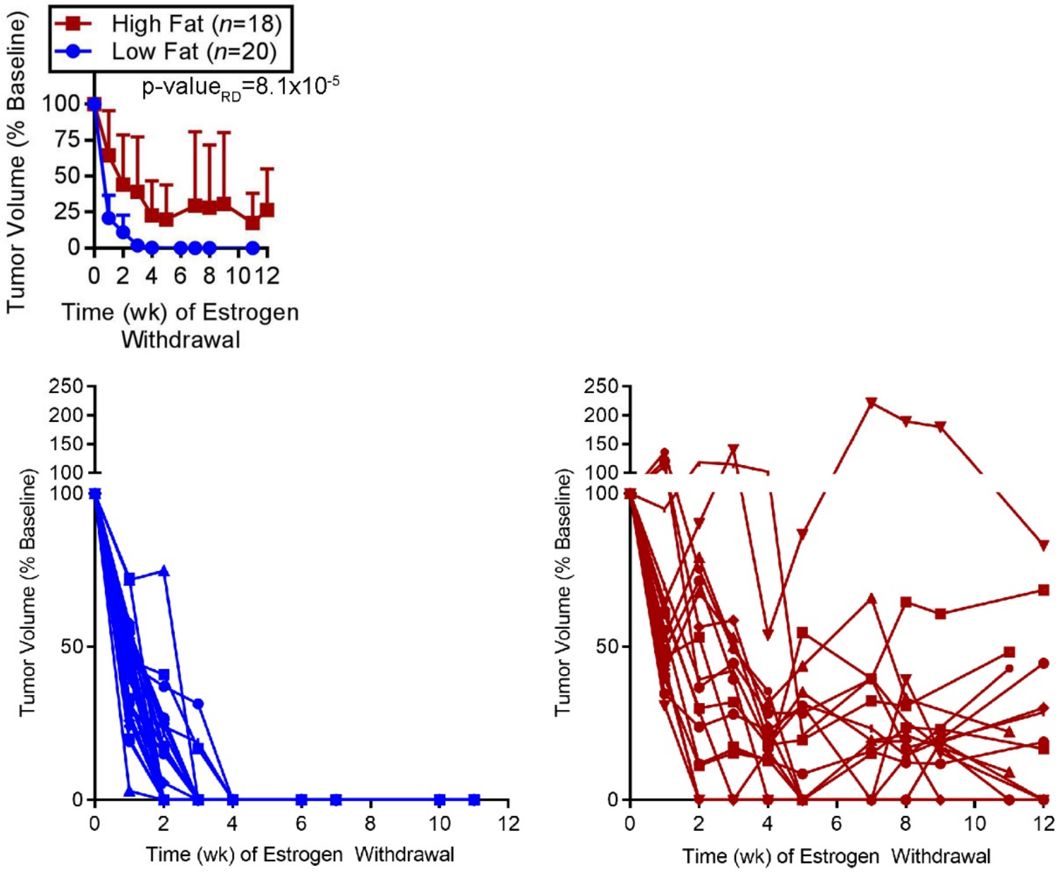

**Figure S19. Immunohistochemical analysis of Ki67, CPT1α, and P-ACC in ER+ breast cancer xenografts treated with either high or low fat diets.** Representative images for Ki67 **(A)**, CPT1α **(B)**, and P-$ACC_{Ser79}$ **(C)** IHC staining of FFPE xenografts are shown. Tumor specimens were harvested from ovx mice on high-fat or low-fat diets following 0 or 90 d of estrogen withdrawal. Summarized data are shown in Fig. 7G-I.

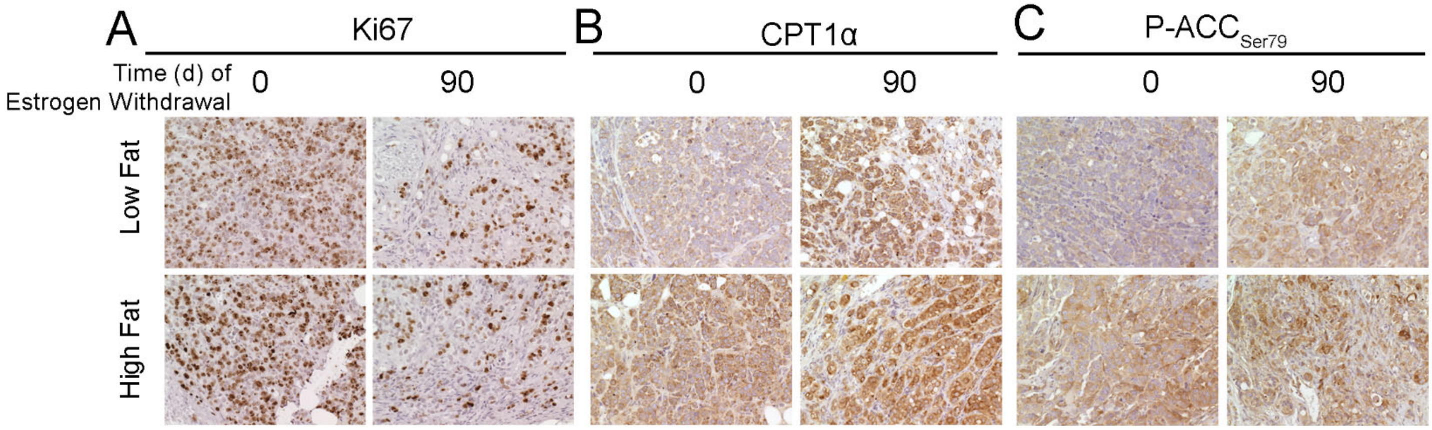

**C. Supplemental Methods**

**Cell culture and RNA interference**

Parental cancer cell lines were obtained from ATCC and used for experiments within 3 months of culture in Dulbecco's Modified Eagle's Medium (DMEM; Corning) with 10% fetal bovine serum (FBS) (Hyclone). For hormone deprivation (HD) experiments, cells were cultured in phenol-red free DMEM (Corning) containing 10% dextran/charcoal-stripped FBS (Hyclone). Controls for HD experiments contained 1 nM E2. Stable cell lines expressing luciferase and GFP were generated using the pLGcmv-CBG-Clover lentiviral vector. The pLGcmv-CBG-Clover lentiviral vector was generated by insertion of the CMV promoter, click beetle green 99 luciferase (Promega) and Clover GFP variant fusion protein separated by a GGGS linker into pFLRu-FH-puro (2). Lentiviral vectors encoding constitutively expressed shRNA targeting AMPKα1/*PRKAA1* (catalog # RHS4430) or non-targeting control (catalog # RHS4346) in the pGIPZ backbone were obtained from Dharmacon. LentiX cells (Clontech) were used to generate lentivirus using standard protocols with pMD2.G and psPAX2 helper plasmids (supplied by D. Trono to Addgene; catalog # 12259 and 12260). Stably transfected target cancer cells were selected for GFP expression by FACS, or for puromycin (1 μg/mL) resistance for 2 wk. siRNA targeting AMPKα1/*PRKAA1* (catalog # SI02622228), AMPKα2/*PRKAA2* (catalog # SI02758595), or non-silencing control (catalog #1027310) were obtained from Qiagen and transfected into cancer cells using Lipofectamine RNAiMax (Thermo Fisher Scientific) per manufacturer's instructions.

**Immunoblotting**

Cells were lysed, and frozen tumors were homogenized and lysed, in RIPA buffer [20 mM Tris, pH 7.4, 150 mM NaCl, 1% NP-40, 10% glycerol, 1 mM EDTA, 1 mM EGTA, 5 mM NaPPi, 50 mM NaF, 10 mM Na β-glycerophosphate, plus fresh Halt protease inhibitor cocktail (Pierce) and 1 mM $Na_3VO_4$ (New England Biolabs)]. Lysates were sonicated, centrifuged at 17,000 x *g* for 10 min at 4°C, and protein in supernatants was quantified using BCA assay (Pierce). Protein extracts were denatured with NuPage (Life Technologies) and reduced with 1.25% β-mercaptoethanol (Sigma). Proteins were heated at 95ºC for 1 min, then separated by SDS-PAGE and transferred to nitrocellulose. Blots were probed with antibodies against AMPKα1, AMPKα2, P-$ACC_{Ser79}$, P-$ULK1_{Ser555}$, vinculin, PGC-1α, CPT1α, actin (Cell Signaling), and total OXPHOS antibody cocktail

(Abcam). HRP-labeled secondary antibodies (GE Healthcare) and ECL substrate (Pierce) were used for signal detection.

**Tumor growth studies**

Female NOD-scid/IL2R$\gamma^{-/-}$ (NSG; NOD.Cg-Prkdcscid Il2rgtm1Wjl/SzJ) mice were obtained from the Norris Cotton Cancer Center Mouse Modeling Shared Resource. Female *Foxn1$^{nu}$/Foxn1$^{nu}$* (nude; JNu) mice were purchased from Jackson Laboratories. Ovariectomized 3-to-4-week-old mice were orthotopically injected bilaterally with MCF-7, HCC-1428, HCC-1500, or MDA-MB-415 luciferase/GFP-expressing cells in PBS, or implanted with fragments of HCI-017 patient-derived xenograft (a gift from Alana Welm, Univ. of Utah). Mice were simultaneously implanted s.c. with a 17β-estradiol (1 mg) beeswax pellet (3). Tumor dimensions were measured twice weekly using calipers (volume = [length$^2$ x width]/2). Once tumors reached ~400 mm$^3$, E2 pellets were removed [*i.e.*, estrogen withdrawal (EW)] and mice were randomized to treatments. Vehicle and metformin were administered via drinking water containing 4% sucrose ± 1 mg/mL metformin. Once tumors completely regressed (*i.e.*, not palpable) following EW, residual tumor burden was monitored by bioluminescence imaging (below).

For drugs tested against dormant residual disease, tumor-bearing mice were treated with EW for 60-90 d to induce dormancy. Baseline bioluminescence was measured on two consecutive days and averaged. Animals were then treated as indicated and bioluminescence was serially measured.

For molecular analysis, tumors were harvested at the indicated time points, fixed in formalin, and paraffin-embedded (FFPE). For dietary studies, isocaloric “high-fat” and “low-fat” diets were obtained from Research Diets, Inc. (catalog # D12451i and D12450Bi) and fed to mice ad libitum.

**Bioluminescence imaging**

Mice were injected i.p. with 100 μL of *in vivo*-grade D-Luciferin (3 mg/mL; Promega) in PBS and placed under isoflurane anesthesia. After a 15-min uptake period, mice were imaged for bioluminescence using a Xenogen IVIS 200 System, and signal values were analyzed using Living Image software (Perkin Elmer). Digital light imaging topography (DLIT) was used to integrate a series of filtered 2-D bioluminescent images with animal surface topography to reconstruct 3-D images.

**Pharmacokinetic analyses**

Female NSG mice were treated metformin (1 mg/mL in drinking water) for 6 wk. Blood was collected retro-orbitally from 3 mice per time point into tubes containing EDTA (0.5 M final concentration) as an anti-coagulant, and centrifuged at 2,000 x *g* for 5 min at 4°C. Plasma was isolated and stored at -80°C. Following blood collection, tumor, liver, and gastrocnemius muscle tissues were then collected, homogenized in 200 μL water, and stored at -80°C.

For metformin measurement in cultured cells, MCF-7 cells were plated and treated as biological triplicates with the indicated concentrations of metformin in growth medium for 16 h. Cells were then rinsed twice with PBS, and scraped into 1 mL water. Scraped cells were frozen at -80°C until analysis.

Metformin concentrations were measured by liquid chromatography with tandem mass spectrometry in technical triplicates, and the average value from technical triplicates for each biological sample was used for calculations. Phenformin was added to all biological samples prior to processing to serve as an internal standard. Samples (50 μL plasma or 10 mg of homogenized tissue) were processed utilizing acetonitrile and Phenomenex iPhree cartridges, and dried under nitrogen at 50ºC. Dried samples were resuspended in water for injection onto an Accucore HILIC 50 mm x 2.1 mm, 2.6 μm column with 10 mm x 2.1 mm HILIC guard cartridge. Isocratic conditions, 25% 100 mM ammonium formate pH 3.2, and 75% acetonitrile over 3 min were utilized at 0.35 mL/min on a Dionex Ultimate 3000 HPLC system. A TSQ Vantage tandem quadrupole mass spectrometer with a HESI-II probe, operated in multiple reaction monitoring mode, was used for positive ion detection, and Xcalibur software was used for data acquisition and processing. MS/MS detection was conducted monitoring 130.09 → 71.100 m/z (collision energy 20 V) for metformin and 206.07 → 60.11 m/z (collision energy 16 V) for phenformin. S-Lens RF amplitudes for metformin and phenformin were 40 V and 60 V respectively. Source parameters were spray voltage 2000 V, vaporizer temperature 275ºC, capillary temperature 350ºC, sheath gas 43, and aux gas 4. Calibration standards were linear over the metformin concentration range from 50-1200 ng/mL (0.39-9.29 μM), with quality controls of 100, 500, and 900 ng/mL (0.77, 3.87, and 6.97 μM). Only calibration standards with >85% accuracy were used for quantitation.

**Serum insulin, blood glucose, and serum free fatty acid analyses**

Mice were fasted overnight, and 200 μL of blood was collected retro-orbitally. Two μL of blood was used to measure glucose concentrations using a glucometer (OneTouch Ultra 2). Remaining blood was allowed to coagulate prior to centrifugation and serum collection. Serum was stored at -80ºC until analysis. Serum insulin and free fatty acid concentrations were measured by ELISA per manufacturers' protocols (Human Insulin ELISA kit, catalog # EZHI-14K from ENDMillipore; Free Fatty Acid Assay Kit, catalog # ab65341 from Abcam).

**Immunohistochemistry and immunofluorescence**

Five-micron sections of FFPE tissue were stained using H&E, immunohistochemistry (IHC), or immunofluorescence (IF). For IHC and IF, sections were deparaffinized in xylene and rehydrated through a graded ethanol series. Heat-induced epitope retrieval was performed using a pressure cooker in either high-pH Tris-EDTA Buffer (pH 9) or low-pH Citrate Buffer (pH 6) (VWR). Sections were then permeabilized with 0.5% Triton X-100 in PBS and treated with 0.3% $H_2 0_2$. Sections were blocked in 5% goat serum for 45 min, then incubated in blocking solution containing primary antibody overnight at 4°C. IHC using antibodies against P-$ACC_{ser79}$, AMPKα2, cleaved caspase 3/7, P-S6 (Cell Signaling), CPT1α (Abcam), ERα (Dako), and Ki67 (Biocare Medical) was performed using Vectastain ABC-HRP kit (Vector Labs). Signal was developed using DAB substrate (Vector Labs). Proportions of positively stained cells in each specimen were counted in 3 random microscopic fields (200x magnification). IHC quantification was performed using HaloVelocity software (Perkin Elmer).

Mitochondrial morphology in xenograft sections was analyzed by IF using the antibody-fluorophore conjugate TOM20-AF594 (Santa Cruz). Images were acquired using a Zeiss LSM 800 with Airyscan microscope using the 63x oil objective and the magnification changer set to 3x zoom. For each Airyscan Z-stack, 35 image slices were taken, with the Z-slices set between 0.16-0.2 μm. IF images were processed and analyzed for mitochondrial count (Number of Connected Components) and length (Average Mitochondrial Length) using MitoGraph software (4). Analysis included ≥30 tumor cells from ≥3 tumors per treatment group.

Mitochondrial membrane potential of cultured cells was measured by incubating cells with tetramethylrhodamine methyl ester (TMRM) for 1 h or 5 h, or with MitoTracker Green (MTG) for 5 h. Staining intensity was measured using the fluorescence image acquisition mode on the IncuCyte S3 Live-Cell Analysis System (Essen Bioscience).

**Tumor cell dissociation and flow cytometry**

GFP-expressing tumor specimens were harvested, rinsed with PBS, and minced. Minced tumor fragments were placed in digestion buffer [(Hank's buffered salt solution (Corning), 7 mg/mL collagenase III (Sigma), 0.2 mg/mL DNAse I (Sigma)] and agitated for 45 min at 37ºC. Tumor digest was passed through a 70-μm filter. P-S6 immunostaining was performed using Foxp3/Transcription Factor Fixation/Permeabilization kit (eBioscience) per manufacturer's protocol with P-S6, CD24, and CD44 antibody-fluorophore conjugates (BD Biosciences). Cells were analyzed using a MACSQuant-10 flow cytometer.

**RNA expression profiling of xenografts**

RNA was isolated from FFPE estrogen-driven tumors (Day 0), acutely EW tumors (Day 3 or 6), and clinically dormant residual tumor cells (Day 82 or 90) using the AllPrep DNA/RNA FFPE Kit (Qiagen). RNA quality was assessed on a fragment analyzer (Advanced Analytical Technologies), and RNA was quantified by Qubit. Nanostring analysis was performed using the PanCancer Pathways Panel (770 genes) on the nCounter FLEX analysis system according to manufacturer's protocol. Data were normalized using nSolver software. The complete normalized dataset is appended in Table S1.

In preparation for RNA-seq, ribo-depleted libraries were prepared from 2.5 μg of total RNA using the GlobinZero Gold (catalog # GZG1206, Illumina) and TruSeq Stranded Total RNA (catalog # RS-122-2201, Illumina) workflows according to manufacturer's instructions. Each library was uniquely barcoded, quantified by qPCR (catalog # KK4824, Kapa Biosystems), and pooled for sequencing on an Illumina NextSeq 500 (2×75-bp). Reads were checked for quality control using FastQC (5), and if necessary were trimmed using

Trimmomatic (6) to trim regions with phred Q >30 (7). High-quality reads were then aligned to reference genome hg19 using STAR (8). Gene counts were normalized by frequency per kilobase million (9). Differential expression of genes was determined using the limma (10) and DESeq2 (11) packages in the R environment (12), and multiple testing correction was performed using the FDR Benjamini-Hochberg method (13). Genes were determined to be significantly differentially expressed if FDR $q \leq 0.05$ and absolute $\log_2$ fold change ≥1. To determine significant differences in the expression of gene signatures between time points, we performed unsupervised sample-wise enrichment analysis of common metabolic and signaling pathways [selected from the Hallmarks (HM), Gene Ontology (GO), Reactome (RM), or Motif gene set(C3) collections] using Gene Set Variation Analysis (GSVA) (14) in R using default arguments with an adjusted p-value significance threshold of 0.2 (1). RNA-seq data are available at NCBI SRA (accession number #PRJNA491455).

**Cellular respiration analysis**

Cells were plated at 20,000-40,000/well in 96-well plates in triplicate at 6 h prior to assay. Oxygen consumption rates (OCR) were serially monitored using the Seahorse XF96 Flux Analyser (Agilent) under the indicated treatment conditions, and after addition of 1 μM oligomycin, 0.5 μM FCCP, and 0.5 μM rotenone.

**Cell growth assays**

Cells were plated at 500-1,000/well in 96-well plates. The following day, cells were washed and medium was changed as indicated. Media and drugs were refreshed every 3-4 d. Growth was serially monitored using the brightfield camera on the IncuCyte S3 Live-Cell Analysis System (Essen Bioscience). Long-term assays (Figs. 4, 5A, and 6B/C) were run for 21-28 d until a well reached 100% confluence. Short-term assays (Figs. 4A and 5B) were run for 7 d.

**Statistical analyses**

Cell growth, IHC, and IF data were analyzed by *t*-test (for two-group experiments), or ANOVA (for experiments with more than two groups) followed by Bonferroni multiple comparison-adjusted posthoc testing between groups. We used the following mathematical model with interpretable parameters to describe a monotonically decreasing pattern to analyze tumor volumes in mice after removal of E2 pellets:

$$V(t) = e^{a_1 + a_2 e^{-(a_3/a_2)t}}, \quad t \geq 0,$$

where $a_1, a_2$, and $a_3$ are positive parameters, and $t$ is time (d). At time $t = 0$ we have $V(0) = e^{a_1+a_2}$, and when $t \to \infty$ we have $V(t) \to e^{a_1}$. On the log scale, the model simplifies and takes the form of the exponential function with asymptote $\ln V(t) = a_1 + a_2 e^{(-a_3/a_2)t}$. Function $V(t)$ is similar to a Gompertz curve that describes tumor growth ($a_2 < 0$) [detailed in (15)].

The *long-term treatment effect* (LT) is measured as the proportion of the tumor volume decrease with respect to the baseline tumor volume, $V(0)$, or in mathematical terms

$$\frac{V(0) - V(\infty)}{V(0)} = \frac{e^{a_1+a_2} - e^{a_1}}{e^{a_1+a_2}} = 1 - e^{-a_2} \simeq a_2.$$

This means that $100a_2\%$ can be interpreted as % tumor reduction with respect to the initial tumor volume. We refer to $a_2$ as the (*relative*) *tumor reduction* parameter that reflects the proportion of tumor decrease on the scale of baseline tumor volume at $t = 0$. For MCF-7 and HCI-017 dormancy models, vehicle-treated tumors regress at a linear rate beyond the limit of caliper measurement; thus, we did not compute long-term treatment effects (when applicable) for those models.

The *rate difference* (RD) is measured as the rate of tumor reduction or growth at time $t = 0$,

$$\frac{1}{V(0)} \frac{dV}{dt}\bigg|_{t=0} = \frac{d\ln V(t)}{dt}\bigg|_{t=0} = \frac{d(a_1 + a_2 e^{-(a_3/a_2)t})}{dt}\bigg|_{t=0} = -a_2(a_3/a_2)e^{-(a_3/a_2)\times 0} = -a_3.$$

This means that $100a_3\%$ can be interpreted as the % tumor volume decrease per unit of time at the time of E2 pellet removal or start of drug treatment. We refer to $a_3$ as the (*relative) rate of tumor reduction* parameter that reflects how fast tumor volume approaches the dormant state (*i.e.*, plateau of volume).

The model in each treatment group was estimated under the assumption that tumor volume at $t = 0$ is mouse-specific, which leads to a nonlinear mixed model:

$$y_{ti} = a_{1i} + a_2 e^{(-a_3/a_2)t} + \varepsilon_{ti}, \quad i = 1,..,n$$

where $n$ is the number of mice in a group, and $a_{1i}$ is a random effect distributed as $N(a_1, \sigma_0^2)$. Standard deviation $\sigma_0$ reflects the heterogeneity of baseline tumor volumes. p-values compare parameters $a_2$ and $a_3$ in treatment groups using the output of nlme function in R (12).

**Analysis of human breast tumor gene expression datasets**

A gene expression signature of response to etomoxir in MCF-7 cells was generated using data downloaded from the Connectivity Map (16) at: https://clue.io/command?q=/sig%20%22etomoxir%22. Among the 992 genes altered in response to treatment with 10 μM etomoxir for 6 or 24 h, 77 genes had a Gene Ontology annotation of "Cell Proliferation" or "Cell Cycle," leaving 915 non-cell cycle-related genes for a subset analysis.

Gene expression profiles of paired human ER+ breast tumor specimens acquired before and after neoadjuvant anti-estrogen therapy were downloaded from the NCBI Gene Expression Omnibus (GEO) for accession numbers GSE71791 (17), GSE20181 (18), and GSE111563 (19). Series matrix files containing $\log_2$-transformed normalized data were collapsed to give one (most variable) probe set per gene using GenePattern. To determine the similarity of a tumor to the etomoxir signature, we generated a *t*-statistic for each tumor in relation to the etomoxir signature as described (20,21). Briefly, we separated etomoxir signature genes based on the direction of their regulation in the etomoxir signature [397 genes were upregulated (z-score ≥ 1) in response to etomoxir; 595 genes were down-regulated (z-score ≤ -1) in response to etomoxir]; for GSE71791, GSE20181, and GSE111563, this included 696, 940, and 691 genes in each respective platform. We then performed a *t*-test of UP genes *vs*. DOWN genes within each tumor. The *t*-statistic was calculated as the two-tailed inverse of the student's *t*-distribution; since etomoxir inhibits FAO, and we wanted to infer FAO activation in tumors, *t*-statistics were multiplied by (-1) for visualization. Thus, a highly positive -(etomoxir *t*-statistic) implies a higher degree of FAO. *t*-statistics were compared between time points by paired *t*-test. Subset analyses were similarly performed using the 915 non-cell cycle-related gene signature of etomoxir response.

**Fatty Acid Oxidation Assay**

Cells were treated with phenol red-free DMEM containing 10% DCC-FBS (hormone-depleted medium, HD) ± 1 nM E2 as indicated. Two to three days prior to assay, cells were trypsinized and reseeded at $10^4$ cells/well in a 96-well plate in HD ± 1 nM E2. Separately, an aliquot of fresh HD medium was mixed with $^3$H-palmitic acid (Perkin Elmer cat # NET043001MC; provided dissolved in ethanol) at a ratio that provided 10 μCi of $^3$H per 100 μL of medium for each well of cells; this mixture was rocked overnight at room temperature to allow the $^3$H-palmitic acid to bind to serum elements. The $^3$H-labeled medium was then aliquoted, and 1 nM E2 and/or 20 μM etomoxir was added. Medium was aspirated from cells, and the $^3$H-labeled medium ± drug(s) was added onto cells (100 μL/well), with 5 biological replicates per treatment condition, and 5 wells containing no cells (to give background readings). Cells were incubated in $^3$H-labeled medium ± drug(s) for 24 h.

$^3$H-labeled medium (~100 μL) was removed from each well and transferred to 1.5-mL centrifuge tubes. Adherent cells were then lysed on ice in 20 uL RIPA buffer containing protease inhibitors; the amount of protein in each well was determined by BCA assay as above. The remaining protocol for processing of $^3$H-labeled medium was adapted from ref. (22). To $^3$H-labeled medium, 100 uL of 10% TCA was added, and tubes were vortexed for 10 sec, incubated at 4ºC for 30 min, and centrifuged at 18 k x *g* for 5 min. Supernatant (120 μL) was transferred to a new centrifuge tube, and 120 μL of 10% TCA was added. Tubes were vortexed, incubated at 4ºC, and centrifuged as above. Supernatant (150 μL) was transferred to a new centrifuge tube, and 300 μL of 2M KCl:HCl solution and 750 μL of methanol:chloroform mixture (2:1 ratio of methanol to chloroform) were added. Tubes were vortexed and centrifuged at 3 k x *g* for 5 min. Supernatant (500 μL) was mixed with 5 mL of scintillation fluid that was analyzed using a liquid scintillation counter.

The scintillation counts-per-minute (CPM) and amount of protein per well from the 5 "no cell" samples were averaged to provide background values, which were then subtracted from each experimental sample. Background-subtracted scintillation values were divided by amount of protein for each well. Data were reported as CPM/mg protein/h, and values relative to E2-treated samples were reported in Fig. 3G.

### D. Supplemental References Cited


1. Subramanian A, Tamayo P, Mootha VK, Mukherjee S, Ebert BL, Gillette MA, *et al.* Gene set enrichment analysis: a knowledge-based approach for interpreting genome-wide expression profiles.

Proceedings of the National Academy of Sciences of the United States of America **2005**;102(43):15545-50.
2. Feng Y, Nie L, Thakur MD, Su Q, Chi Z, Zhao Y*, et al.* A multifunctional lentiviral-based gene knockdown with concurrent rescue that controls for off-target effects of RNAi. Genomics, proteomics & bioinformatics **2010**;8(4):238-45.
3. DeRose YS, Gligorich KM, Wang G, Georgelas A, Bowman P, Courdy SJ*, et al.* Patient-derived models of human breast cancer: protocols for in vitro and in vivo applications in tumor biology and translational medicine. Current protocols in pharmacology / editorial board, SJ Enna **2013**;Chapter 14:Unit14 23.
4. Harwig MC, Viana MP, Egner JM, Harwig JJ, Widlansky ME, Rafelski SM*, et al.* Methods for imaging mammalian mitochondrial morphology: A prospective on MitoGraph. Analytical biochemistry **2018**;552:81-99.
5. Andrews S. FastQC: a quality control tool for high throughput sequence data. 2010.
6. Bolger AM, Lohse M, Usadel B. Trimmomatic: a flexible trimmer for Illumina sequence data. Bioinformatics **2014**;30(15):2114-20.
7. Ewing B, Hillier L, Wendl MC, Green P. Base-calling of automated sequencer traces using phred. I. Accuracy assessment. Genome research **1998**;8(3):175-85.
8. Dobin A, Davis CA, Schlesinger F, Drenkow J, Zaleski C, Jha S*, et al.* STAR: ultrafast universal RNA-seq aligner. Bioinformatics **2013**;29(1):15-21.
9. Garber M, Grabherr MG, Guttman M, Trapnell C. Computational methods for transcriptome annotation and quantification using RNA-seq. Nature methods **2011**;8(6):469-77.
10. Ritchie ME, Phipson B, Wu D, Hu Y, Law CW, Shi W*, et al.* limma powers differential expression analyses for RNA-sequencing and microarray studies. Nucleic acids research **2015**;43(7):e47.
11. Love MI, Huber W, Anders S. Moderated estimation of fold change and dispersion for RNA-seq data with DESeq2. Genome biology **2014**;15(12):550.
12. R Core Team. R: A language and environment for statistical computing. . Vienna, Austria: R Foundation for Statistical Computing; 2017.
13. Benjamini Y, Hochberg Y. Controlling the False Discovery Rate - a Practical and Powerful Approach to Multiple Testing. J Roy Stat Soc B Met **1995**;57(1):289-300.
14. Hanzelmann S, Castelo R, Guinney J. GSVA: gene set variation analysis for microarray and RNA-seq data. BMC bioinformatics **2013**;14:7.
15. Demidenko E. Mixed Models: Theory and Applications with R. **2013**.
16. Subramanian A, Narayan R, Corsello SM, Peck DD, Natoli TE, Lu X*, et al.* A Next Generation Connectivity Map: L1000 Platform and the First 1,000,000 Profiles. Cell **2017**;171(6):1437-52 e17.
17. Patani N, Dunbier AK, Anderson H, Ghazoui Z, Ribas R, Anderson E*, et al.* Differences in the transcriptional response to fulvestrant and estrogen deprivation in ER-positive breast cancer. Clin Cancer Res **2014**;20(15):3962-73.
18. Miller WR, Larionov A, Anderson TJ, Evans DB, Dixon JM. Sequential changes in gene expression profiles in breast cancers during treatment with the aromatase inhibitor, letrozole. Pharmacogenomics J **2012**;12(1):10-21.
19. Selli C, Turnbull AK, Pearce DA, Li A, Fernando A, Wills J*, et al.* Molecular changes during extended neoadjuvant letrozole treatment of breast cancer: distinguishing acquired resistance from dormant tumours. Breast Cancer Res **2019**;21(1):2.
20. Gibbons DL, Lin W, Creighton CJ, Zheng S, Berel D, Yang Y*, et al.* Expression signatures of metastatic capacity in a genetic mouse model of lung adenocarcinoma. PLoS One **2009**;4(4):e5401.
21. Creighton CJ, Casa A, Lazard Z, Huang S, Tsimelzon A, Hilsenbeck SG*, et al.* Insulin-like growth factor-I activates gene transcription programs strongly associated with poor breast cancer prognosis. J Clin Oncol **2008**;26(25):4078-85.
22. Dunning KR, Cashman K, Russell DL, Thompson JG, Norman RJ, Robker RL. Beta-oxidation is essential for mouse oocyte developmental competence and early embryo development. Biol Reprod **2010**;83(6):909-18.
